# Examining the boundary sharpness coefficient as an index of cortical microstructure and its relationship to age and sex in autism spectrum disorder

**DOI:** 10.1101/2020.07.09.196212

**Authors:** Emily Olafson, Saashi Bedford, Gabriel A. Devenyi, Raihaan Patel, Stephanie Tullo, Min Tae M. Park, Olivier Parent, Evdokia Anagnostou, Simon Baron-Cohen, Edward T. Bullmore, Lindsay R. Chura, Michael C. Craig, Christine Ecker, Dorothea L. Floris, Rosemary J. Holt, Rhoshel Lenroot, Jason P. Lerch, Michael V. Lombardo, Declan G. M. Murphy, Armin Raznahan, Amber N. V. Ruigrok, Michael D. Spencer, John Suckling, Margot J. Taylor, MRC AIMS Consortium, Meng-Chuan Lai, M. Mallar Chakravarty

## Abstract

Autism spectrum disorder (ASD) is associated with atypical brain development. However, the phenotype of regionally specific increased cortical thickness observed in ASD may be driven by several independent biological processes that influence the gray/white matter boundary, such as synaptic pruning, myelination, or atypical migration. Here, we propose to use the boundary sharpness coefficient (BSC), a proxy for alterations in microstructure at the cortical gray/white matter boundary, to investigate brain differences in individuals with ASD, including factors that may influence ASD-related heterogeneity (age, sex, and intelligence quotient). Using a vertex-based meta-analysis and a large multi-center magnetic resonance structural imaging (MRI) dataset, with a total of 1136 individuals, 415 with ASD (112 female; 303 male) and 721 controls (283 female; 438 male), we observed that individuals with ASD had significantly greater BSC in the bilateral superior temporal gyrus and left inferior frontal gyrus indicating an abrupt transition (high contrast) between white matter and cortical intensities. Increases were observed in different brain regions in males and females, with larger effect sizes in females. Individuals with ASD under 18 had significantly greater BSC in the bilateral superior temporal gyrus and right postcentral gyrus; individuals with ASD over 18 had significantly increased BSC in the bilateral precuneus and superior temporal gyrus. BSC correlated with ADOS-2 CSS in individuals with ASD in the right medial temporal pole. Importantly, there was a significant spatial overlap between maps of the effect of diagnosis on BSC when compared to cortical thickness. These results invite studies to use BSC as a possible new measure of cortical development in ASD and to further examine the microstructural underpinnings of BSC-related differences and their impact on measures of cortical morphology.

## Introduction

Neuroimaging studies of autism spectrum disorder (ASD) have repeatedly observed early increases in cortical thickness (Bedford et al. 2019; Anagnostou and Taylor 2011; Hazlett et al. 2017; Courchesne et al. 2011; Park et al. 2018) and altered structural and functional connectivity in individuals with ASD, particularly early in life (Hahamy, Behrmann, and Malach 2015; Rudie et al. 2012; Just et al. 2012). The early developmental period, which is coincident with ASD-onset, is a particularly sensitive time with respect to neuronal migration and cortical myelination. Indeed, neuronal migration disruptions from the ventricular and subventricular zones during early neocortex development have been observed in ASD (Pinto et al. 2014; Huguet, Ey, and Bourgeron 2013; Reiner et al. 2016), potentially leading to the presence of supernumerary neurons proximal to the gray/white matter boundary. Additionally, maturational abnormalities of intracortical myelin, a key cortical maturational feature (Grydeland et al. 2013; Deoni et al. 2015), have been associated with ASD (Graciarena et al. 2018; Zikopoulos and Barbas 2010; Canali et al. 2018). This developmental variation may underlie measures seeking to capture so-called ‘blurring’ at the interface of the cortex and the underlying superficial white matter, as measured using structural magnetic resonance imaging (MRI) (Norbom et al. 2019; Mann et al. 2018; Andrews et al. 2017; Bezgin, Lewis, and Evans 2018). Despite advances in ASD research generated by studying the ratio of the tissue intensities between these two compartments, there are limitations that may complicate the interpretation of this measure as an index of cortical microstructure. Firstly, the ratio of intensities between the tissue classes is dependent on the placement of the boundary that separates them. However, the boundary placement may be influenced by signal ambiguity that arises from a less defined gray/white matter boundary, thereby potentially confounding the measurement of interest.

Further, histological studies have shown that more convex regions such as gyral crowns are more myelinated in deeper cortical layers compared to concave regions, such as sulcal folds (Sereno et al. 2013; Bok 1959); thus measures of cortical microstructure are negatively correlated with cortical curvature, as T1-relaxation times are observed to be higher in gyral crowns and lower in sulcal folds (Sereno et al. 2013; Waehnert et al. 2016). In ASD, microstructural measurements obtained using the standard ratio of tissue intensities may be confounded by the well-characterized group differences in gyrification and curvature (Libero et al. 2019; Kohli et al. 2018).

Here, we propose to partly overcome these limitations using our newly-developed metric, boundary sharpness coefficient (BSC), defined as the growth rate parameter of a sigmoid curve fit to the cortical intensity profile running perpendicular to the boundary surface (inspired by (Avino and Hutsler 2010)). Using this new measure that potentially reflects perturbations to neuronal migration and/or intracortical myelination, we performed a large-scale analysis using images collected from multiple acquisition sites and analyzed them with a meta-analytic technique (Bedford et al. 2019). In light of previous findings from our group (Bedford et al. 2019) and others (Anagnostou and Taylor 2011; Hazlett et al. 2017; Courchesne et al. 2011; Schuetze et al. 2016) of increased cortical thickness in ASD, we sought to further examine if previous studies of cortical thickness increases in ASD may have been influenced by cortical boundary abnormalities.

Given results from previous neuroimaging and histological studies (Avino and Hutsler 2010; Mann et al. 2018; Bezgin, Lewis, and Evans 2018; Andrews et al. 2017), we expected to find a greater degree of cortical blurring in individuals with ASD relative to typically developing individuals, or a decrease in BSC. Since T1w intensity is thought to reflect underlying myeloarchitecture more so than cytoarchitecture (Eickhoff et al. 2005), blurring may arise from differences intracortical myelination: an increase in myelin content (and of T1w signal) in the lower layers of the cortex would appear in T1w MRI as a more blurred transition in intensity moving from gray to white matter. An alternative process that may increase cortical blurring in ASD is neuronal migration. Neuronal migration defects are predicted to result in supernumerary neurons in the white matter compartment directly below the cortex (Andrews et al. 2017; Avino and Hutsler 2010; Chun and Shatz 1989), which would also result in a greater degree of blurring captured with BSC.

Another goal of this study was to examine the influence of clinical heterogeneity on BSC. Based on previous studies of cortical anatomy, we expected age-related differences in BSC to be greatest in younger ASD individuals (Khundrakpam et al. 2017; Bedford et al. 2019), to vary by sex (Irimia et al. 2017; Zeestraten et al. 2017; Greenberg et al. 2018; M.-C. Lai et al. 2015; M. C. Lai et al. 2017) and full-scale IQ (FIQ) (Bedford et al. 2019; Lotspeich et al. 2004).

## Methods

### Study participants

Data included here were acquired from previous studies by the Hospital for Sick Children (Canada), the Cambridge Family Study of Autism (UK) and the UK Medical Research Council Autism Imaging Multicentre Study (UK MRC AIMS). We also included publicly available data from the Autism Brain Imaging Data Exchange (ABIDE) I and II datasets (Adriana Di Martino et al. 2017; A. Di Martino et al. 2014) (Table 1 and 2). All data used in this study was preprocessed by SB for analysis in (Bedford et al. 2019), with an original sample size of 3145 participants (1415 individuals with ASD (1165 male/250 female) and 1730 controls (1172 male/558 female), aged 2-65 years. Sample characteristics (including image processing, and which subjects were excluded due to poor motion or segmentation quality) are equivalent between studies, with the exception of the NIMH site, which was not included in this study because the partial volume effects from its lower resolution would compromise the accuracy of BSC measurement. Quality control methodology is outlined in detail in a recent manuscript from our group (Bedford et al. 2019).

**Table 1.**
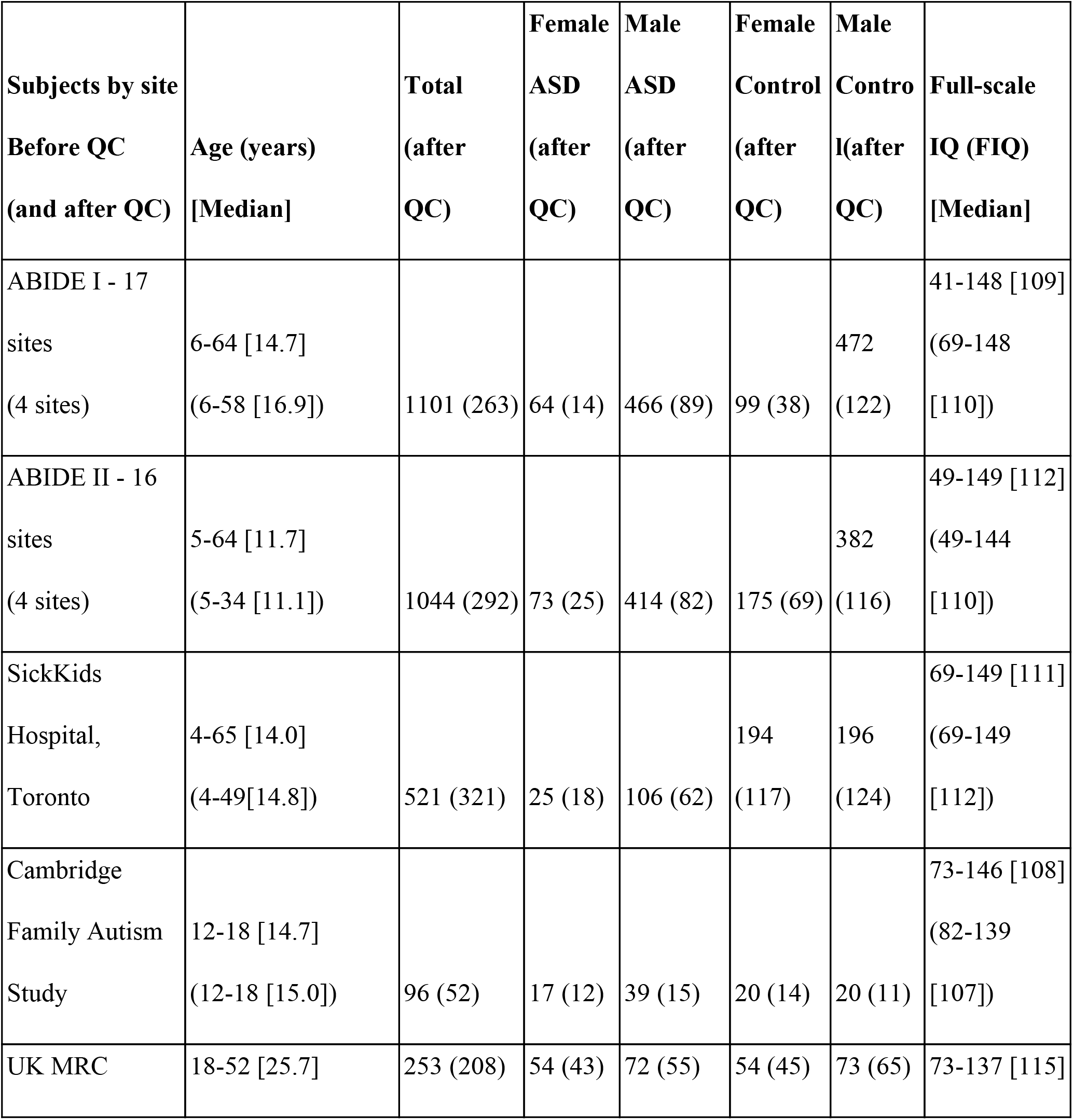
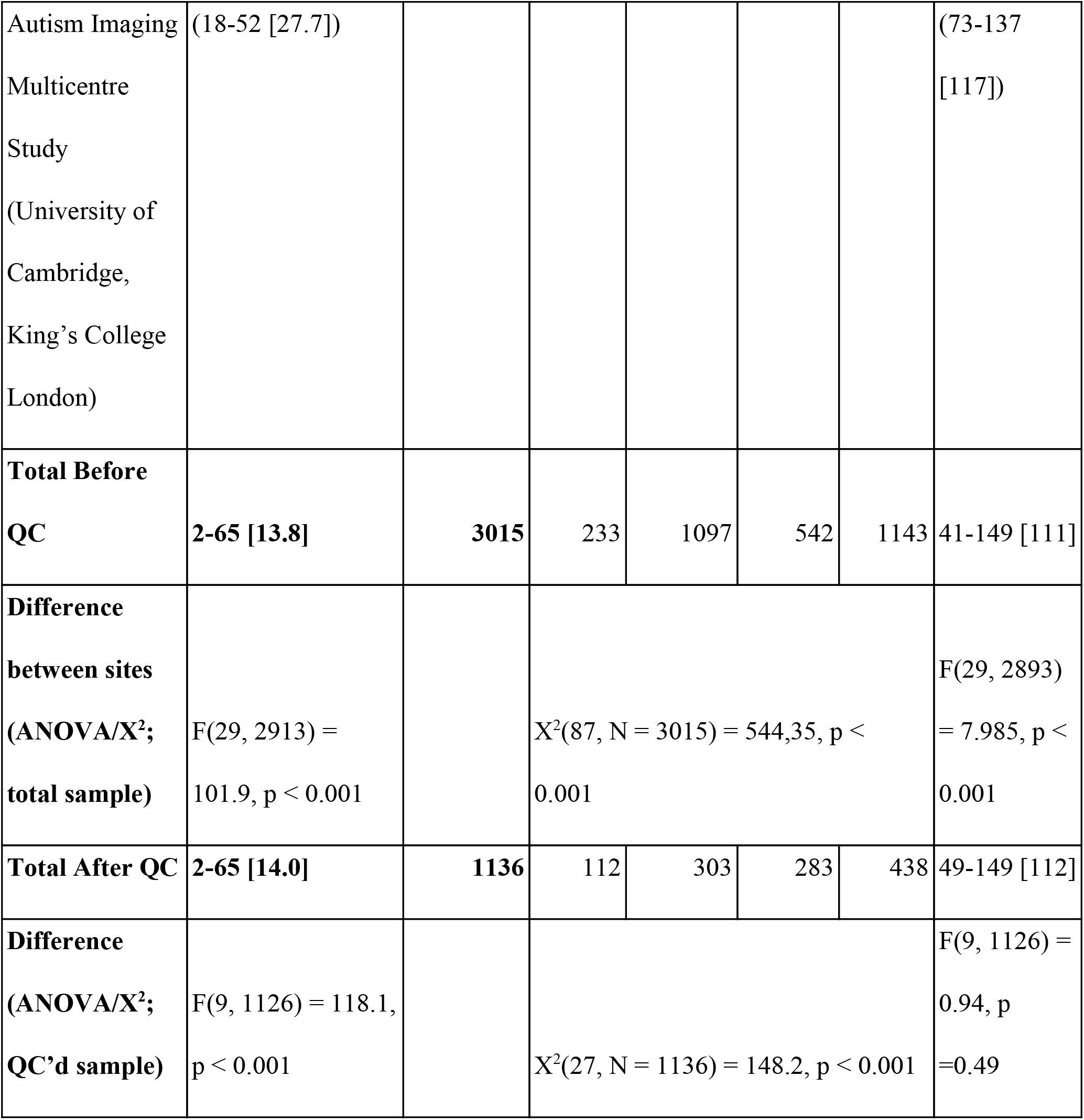
Subject demographics of the cohort under study. Group differences in age, sex distribution, and FIQ were also examined using t-tests (for continuous variables) and Chi-squared tests (for categorical variables).

**Table 2.** Subject demographics broken down by ABIDE I and II site.

| <u>Subjects by ABIDE</u> | Age | Total | F- ASD | M- ASD | F- Ctl | M- Ctl | Full-scale |
| --- | --- | --- | --- | --- | --- | --- | --- |
| <u>site Before QC (and</u> | (years) | (after QC) | (after | (after | (after | (after | IQ (FIQ) |

| **<u>after QC)</u>** | **[Median]** |  | **QC)** | **QC)** | **QC)** | **QC)** | **[Median]** |
| --- | --- | --- | --- | --- | --- | --- | --- |
| Kennedy Krieger<br>Institute<br>(KKI; ABIDE I&II) | 8-13<br>[10.3]<br>(8-13<br>[10.4]) | 256<br>(148) | 19<br>(11) | 58<br>(21) | 65<br>(45) | 123<br>(71) | 63-143<br>[113]<br>(69-143<br>[114]) |
| Ludwig Maximilians<br>University Munich<br>(MAX MUN;<br>ABIDE I) | 7-58<br>[26.0]<br>(7-58<br>[29.5]) | 57<br>(38) | 3<br>(3) | 21<br>(14) | 4<br>(4) | 29<br>(17) | 79-133<br>[112]<br>(93-133<br>[112]) |
| New York University<br>(NYU; ABIDE I &<br>II) | 5-39<br>[10.9]<br>(5-39<br>[12.1]) | 289<br>(179) | 19<br>(10) | 135<br>(74) | 28<br>(19) | 107<br>(76) | 67-148<br>[109]<br>(67-148<br>[109]) |
| Oregon Health and<br>Science University<br>(OHSU; ABIDE I &<br>II) | 7-15<br>[11.0]<br>(8-15<br>[11.0]) | 121<br>(97) | 7<br>(5) | 43<br>(33) | 29<br>(25) | 42<br>(34) | 69-140<br>[115]<br>(69-136<br>[114]) |
| San Diego State<br>University<br>(SDSU; ABIDE I &<br>II) | 7-18<br>[13.8]<br>(8-18<br>[14.4]) | 94<br>(45) | 8<br>(6) | 39<br>(16) | 8<br>(3) | 39<br>(20) | 66-141<br>[105]<br>(66-139<br>[104]) |
| University of<br>Michigan<br>(UM; ABIDE I) | 8-28<br>[14.0]<br>(9-28<br>[15.8]) | 145<br>(55) | 10<br>(4) | 58<br>(17) | 18<br>(12) | 59<br>(22) | 76-147<br>[108]<br>(78-147)<br>[110]) |

Supplementary materials of that work (Supplementary Methods: 2. Quality control (QC) and site elimination; 3. Image processing; 7. Quality control analysis) provide detailed accounts of our quality control choices for both the raw and processed data. In total, after segmentation quality control and motion quality control, there remained 1136 subjects: 415 with ASD (303 male/112 female) and 721 controls (438 male/283 female).

### Image processing

Data was preprocessed using the minc-bpipe-library pipeline, including N4 bias field inhomogeneity correction (Tustison et al. 2010). Data was then processed through the CIVET 1.1.12 cortical segmentation and cortical thickness pipeline which generated gray/white and pial surfaces and transformed subject brain volumes into standard MNI space (Ad-Dab’bagh Y. 2006). All intensity sampling and surface generation was performed in standard space.

### BSC calculation

For each of the 77,212 vertices (which exclude all vertices on the midline wall) on the cortical surface of each subject, signal intensity was measured at 10 surfaces that span the region around the gray/white boundary. Together, the 10 intensity samples derived from these new surfaces cover the bottom quarter of the cortex (the quarter that interfaces with the gray/white boundary) and a small portion of white matter below the gray/white boundary (Fig. 1).

**Figure 1.**
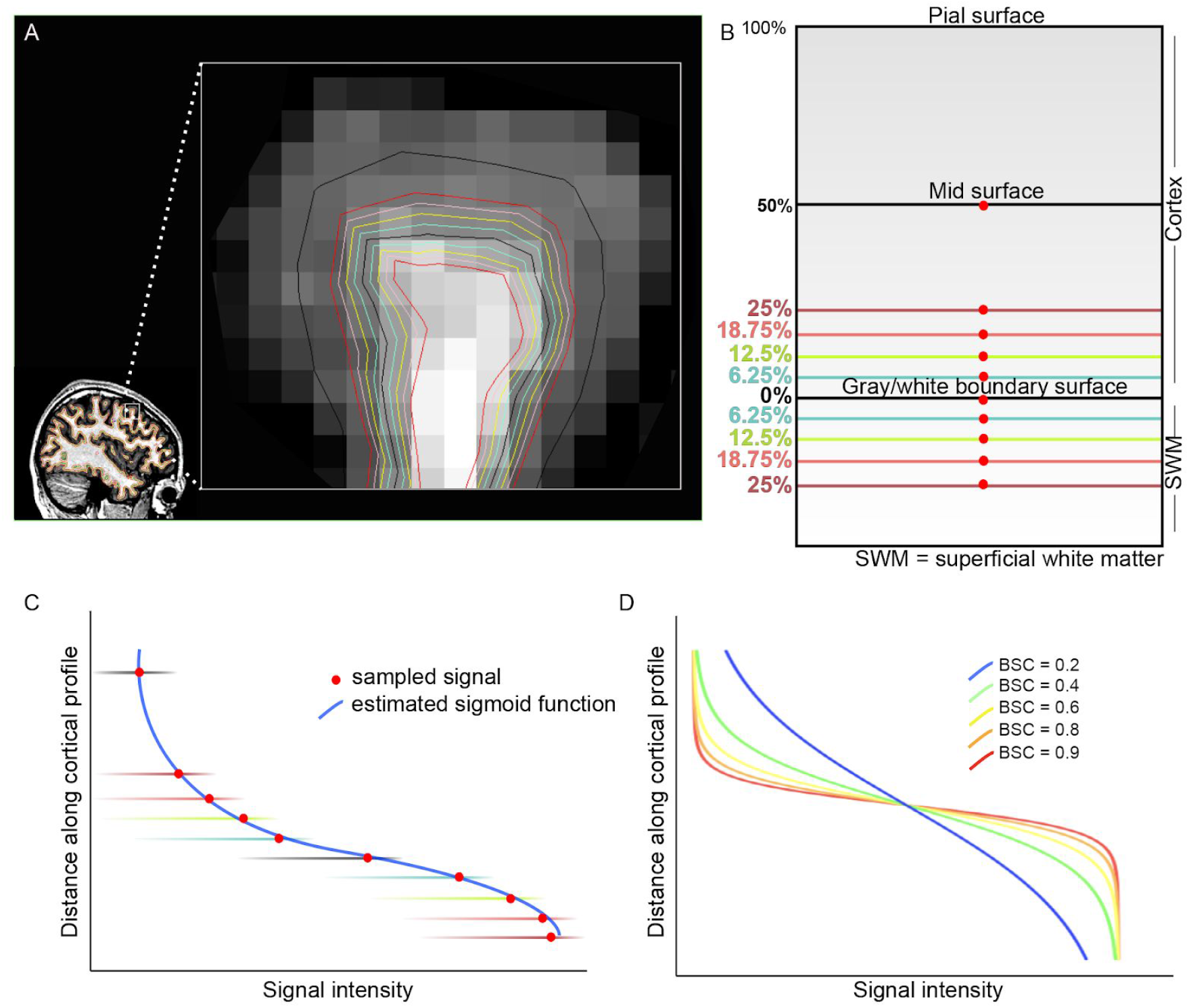
Intensity sampling and sigmoid fit method used to calculate BSC for each vertex. At each vertex, the T1w image intensity was measured at 10 cortical surfaces surrounding the gray/white boundary, including the mid-surface, gray/white boundary surface, and a total of 8 newly-generated gray and white surfaces equally spaced apart (A, B). A sigmoid curve (Equation 1) was fit to the 10 sample points (C) and parameter *BSC*, reflecting the sigmoid growth rate (D), was log-transformed to create the measure that is the BSC at that vertex. Higher BSC values reflect a quicker transition between gray and white matter and a less blurred cortical boundary, whereas lower BSC values reflect a slower transition between gray and white matter and a more blurred cortical boundary (D). The tissue contrast ratio was calculated by dividing the intensity sampled at the gray 25% surface by the intensity sampled at the white 25% surface.

Gray matter surfaces were created at increasing percentile fractions of the cortical thickness (0%, 6.25%, 12.5%, 18.75%, 25%, 50%), from the boundary to the pial surface (Fig. 1a,b). New white matter surfaces were generated at the same distance from the gray/white boundary surface as the gray matter surfaces, but in the direction of the white matter, with the exception of the white matter equivalent of the 50% gray matter surface, which was not included because it crossed into neighboring cortex in thin gyral crowns at certain vertices. See Supplementary Methods section S2 “Surface generation” for more details.

A sigmoid curve (Equation 1) was then fit to the 10 sample points using a non-linear least squares estimator (Fig. 1c); see Supplementary Methods section S3 “Model fit” for more details.

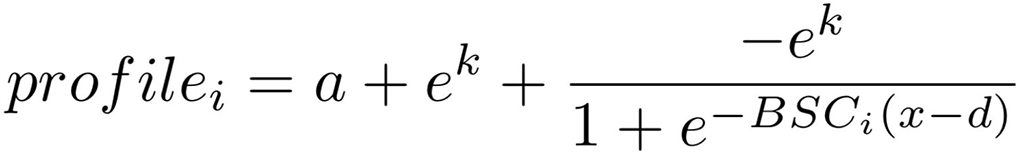

**Equation 1**. Sampled intensities at vertex *i* (*profile_i_*) and their distance along the axis perpendicular to the gray matter boundary surface were input to a nonlinear least-squares estimator to fit a sigmoid function. The *d* parameter reflects the translation of the sigmoid along the x-axis, where x is a vector of the displacement along the axis that runs orthogonal to the boundary, increasing towards the pial surface, parameter *a* translates the sigmoid along the T1 intensity, parameter *k* reflects the height of the sigmoid, and parameter *BSCi* reflects the growth rate (steepness) of the sigmoid.

BSC at each vertex (*BSC* parameter of the sigmoid curve) was extracted from the estimated sigmoid curve (Fig. 1d), where a higher BSC reflects greater boundary “sharpness”, or a faster transition in intensity moving from gray matter to white matter (effectively a measure of contrast at the gray/white matter surface, with higher BSC reflecting high contrast which we take to mean a bigger difference between the microstructural composition of tissue compartments). See Supplementary Fig. S5 for representative sigmoid fits and associated BSC parameters, and Supplementary Fig. S6 for a spatial distribution of model convergence failures prior to spatial smoothing. These values were then log-transformed and smoothed with a 20 mm full-width half-max (FWHM) smoothing kernel to reduce the effect of noise and simulate a Gaussian distribution. No smoothing was applied to the sampled intensity values prior to the model fit. Smoothing kernels at 10mm (Norbom et al. 2019; Mann et al. 2018), 20mm (Bezgin, Lewis, and Evans 2018), and 30mm (Salat et al. 2009; Uribe et al. 2018) FWHM have been used in prior studies investigating the tissue contrast ratio. Here, a 20mm kernel was used with the a priori hypothesis that effects would be present at this intermediate kernel width. The analysis of main effect was repeated with a smoothing kernel of 10mm FWHM. Values were then resampled to a common surface mesh to enable cross-subject comparisons (Lerch and Evans 2005).

By measuring T1 signal at multiple distances on either side of the gray/white boundary and fitting an adequately flexible sigmoid curve to the measurements, BSC is theoretically able to capture cortical blurring wherever it occurs relative to the segmented boundary and thus avoids the circular boundary placement problem mentioned in the Introduction. Finally, given that BSC maps also correlated with mean curvature (Supplementary Fig. S2b) in certain parts of the cortex, BSC values were further residualized for mean curvature at the vertex-wise level across all subjects within each site in standard space (Sereno et al. 2013).

### Relationship between in-scanner motion and BSC

In order to assess the potential impact of motion on data that passed quality control, we obtained motion parameters from fMRI data in 2 sites (ABIDE I KKI; ABIDE II OHSU) which include subjects aged 7-15, a demographic for whom motion is traditionally higher overall. Though an indirect measure of T1 motion, fMRI realignment parameters acquired during the same session have been shown to correlate with anatomical scan motion (A. Alexander-Bloch et al. 2016). Linear regression was performed at each lobe (frontal, temporal, parietal, occipital, cingulate) to estimate the effect of in-scanner motion on BSC (Supplementary Methods S7).

### Code availability

Code for all processing steps, including surface generation and BSC calculation is publicly available on the CoBrALab GitHub: https://github.com/CoBrALab/BSC.

### Statistical Analysis of BSC

Data was amassed from multiple sites in order to be sufficiently powered to analyse sources of heterogeneity in ASD. However, analysing multiple sites presents the challenge of site-specific confounds, such as scanner model, scan acquisition protocols, and sample demographics. These limitations were addressed by the prospective meta-analysis technique, where each site was treated as a separate study, and the results were pooled across sites to determine significance at each vertex (Thompson et al. 2014; van Erp et al. 2016; Bedford et al. 2019).

Meta-analyses were performed at a vertex-wise level (i.e., performing regressions at each vertex across the brain). Specifically, within each site, for each vertex across the brain where BSC was estimated, a linear regression model was performed to derive per-site Cohen’s d effect sizes for the main effect of each variable of interest. Then, the final effect size representing the contribution of all sites was calculated in a random-effects meta-analysis (Borenstein et al. 2010) using the *metafor* package in R 3.4.0 (https://www.r-project.org/). See Supplementary Methods section S5 “Statistical Models” for more details. How models were chosen to study the impact of age, sex, and IQ are further detailed below. Final p-values were adjusted for multiple comparisons across all tests performed (all vertices of all analyses performed; as previously in (Bedford et al. 2019) using the False Discovery Rate (FDR) correction, which controls the proportion of null hypotheses that are falsely rejected (Genovese, Lazar, and Nichols 2002).

### Akaike Information Criterion (AIC) analysis of variable importance

The importance of age (linear term), age (quadratic term), sex, and FIQ were examined by using a vertex-wise Akaike Information Criteria (AIC) analysis. The AIC is a measure that assesses the relative quality of statistical models, where a model with the lowest AIC is considered the best fit for the data (Mazerolle 2006). AIC takes into account both accuracy and parsimony, because it carries a penalty for increasing the number of free parameters in the model. Within each site, the AIC was calculated at each vertex for the linear regression model without the variable of interest (e.g. diagnosis only), with the variable of interest (e.g. age + diagnosis) and with its interaction with diagnosis (e.g. age + diagnosis + age*diagnosis). The percentage of sites for which each of the above models was the best-fitting model, according to AICc (for small sample sizes) was calculated for each vertex as a weighted average based on site size (number of subjects scanned at each site after QC).

Supplementary Fig. S7 displays these results for age, Supplementary Fig. S8 for sex, and Supplementary Fig. S9 for FIQ.

Based on the comparison of the models using AIC, it was determined that age (but not sex or FIQ) was an important explanatory variable at a substantial proportion of vertices across the brain for BSC, thus motivating our investigation into how this factor influences BSC in ASD. See Supplementary Methods section S5 “Statistical Models” for more details.

### Age-focused analyses

The impact of age was examined by stratifying subjects by age, performing separate meta-analyses in subjects who were 18 years old and below, and in subjects who were above 18 years old, including sex and FIQ as covariates. Additionally, an age-centered analysis was performed to assess the trajectory of group differences by shifting the age at which group differences are assessed by 4-year intervals. This type of analysis allows for the interpretation of group differences at various 4-year cross-sections without splitting the dataset into age ranges, maximizing statistical power. The Cohen’s d effect size for the main effect of diagnosis was calculated for each site in each model (one for each age interval) and pooled in a random-effects meta-analysis in the same manner as the case-control comparisons.

### Associations between BSC and ASD symptoms/characteristics

Since consistent measures of ASD symptoms or characteristics were not available across all sites, analyses were performed on a subset of individuals who had the same clinical measures. We chose the measure which had the largest number of individuals available, which was the ADOS-2 Calibrated Severity Scores (CSS) to examine overall symptom severity (N =172; also conducted separately in males [N =139] and females [N =33]). The analysis of the relationship between BSC and severity measures (as measured by ADOS-2 CSS) was performed by conducting a multiple regression analysis, calculating the semi-partial correlation of the ADOS-2 scores with BSC at each vertex, per site, with age, sex, and FIQ included in the model, in individuals with ASD only. The semi-partial correlation was then pooled across sites in a random effects meta-analysis, in the same fashion as the Cohen’s d effect size in a typical meta-analysis (Bedford et al. 2019).

### Assessing spatial overlap between BSC and cortical thickness maps

The spatial correspondence between maps of the effect size of diagnosis on BSC and the effect size of diagnosis on cortical thickness was determined using a ‘spin test’ (A. F. Alexander-Bloch et al. 2018), which generates null estimates of overlap by applying random rotations to spherical projections of a cortical surface. The Pearson’s correlation coefficient between the two original spatial maps is compared to the Pearson’s correlation coefficient measured between one original spatial map and the other map’s rotated permutations. Maps of cortical thickness were developed in (Bedford et al. 2019) from a near-identical dataset (excluding the NIMH dataset here).

## Results

### The tissue intensity ratio correlates with mean curvature

To determine the extent to which tissue intensity ratio may be influenced by cortical curvature, we assessed the correlation between mean curvature and the tissue intensity ratio (see Supplementary Methods section S4 “Tissue intensity ratio” for methods). In a vertex-wise single-subject analysis, we found that the tissue intensity ratio was lower in gyri compared to sulci (Supplementary Fig. S1). Furthermore, in a cross-subject analysis to determine in which areas of the brain this relationship is present, we found a negative correlation between the tissue intensity ratio and mean curvature for most vertices in the brain (Supplementary Fig. S2a, c, d), including vast areas of the frontal, parietal, and occipital lobes. These results are consistent with the hypothesis that the impact of curvature on cortical T1w intensity extends to the tissue intensity ratio, and calls for a measure of cortical contrast between the gray and white matter that is less susceptible to bias by other cortical features. When assessed for its relationship to curvature, we found that BSC correlated with mean curvature in regions of the anterior frontal, parietal, and temporal lobes (Supplementary Fig. S2b), and did not correlate in any regions after regressing the values against mean curvature at the vertex level.

### Average BSC map recapitulates T1/T2 ratio map

We observed regional differences in mean BSC across healthy individuals ages 22-35 within the Sick Kids (Toronto) cohort, with the lowest mean BSC in areas such as the motor-somatosensory strip in the central sulcus, the visual cortex in the occipital lobe, and in early auditory areas in the Sylvian fissure, matching areas that were reported to contain the heaviest intracortical myelination as measured by the T1/T2 ratio in Glasser et al., (2011).

### Normative effect of age and sex on BSC in control subjects

In order to contextualize diagnostic group differences in BSC, we performed a vertexwise analysis of the main effects of age, age squared, and sex on BSC across all control individuals in the Sick Kids (Toronto) cohort (165F/149M; aged 4-65) (Supplementary methods section S6). Main effects of age on BSC were significant across the cortex with a negative relationship in association cortices and positive relationship in sensory areas. Likewise, age-squared was negatively related to BSC in association cortices (an inverted U-shaped relationship), and positively related in primary sensory areas (U-shaped relationship) (Supplementary Figure S3a, b). Significant sex differences were only observed in the bilateral medial temporal cortices where males have higher BSC than females (Supplementary Figure S3c).

### Motion does not correlate with BSC

Multiple regression between average BSC and in-scanner motion (as estimated by Framewise Displacement; see Supplementary Methods S7) did not provide evidence for a relationship between BSC and in-scanner motion (Supplementary Figure S4; Table 3).

**Table 3.** Results from linear regressions between average BSC in various compartments (rows) and average Framewise Displacement, a measure of in-scanner motion.

(a) ABIDE I KKI
|  | Estimate (B_1) | t-statistic | p-value (FDR corrected) |
| --- | --- | --- | --- |
| **Whole brain avg. FD** | 0.0091022 | 0.80038 | 0.6848 |
| **Frontal lobe avg. FD** | 0.0082202 | 0.57611 | 0.6848 |
| **Parietal lobe avg. FD** | 0.013204 | 1.2376 | 0.6848 |
| **Temporal lobe avg. FD** | 0.0091022 | 0.80038 | 0.6848 |
| **Occipital lobe avg. FD** | 0.0059099 | 0.58677 | 0.6848 |
| **Cingulate/insula avg. FD** | 0.001678 | 0.089094 | 0.9298 |

(b) ABIDE II OHSU
|  | Estimate (B_1) | t-statistic | p-value (FDR corrected) |
| --- | --- | --- | --- |
| **Whole brain avg. FD** | 0.0081261 | 0.84114 | 0.6226 |
| **Frontal lobe avg. FD** | 0.008641 | 0.77198 | 0.6226 |
| **Parietal lobe avg. FD** | 0.0071437 | 0.69398 | 0.6226 |
| **Temporal lobe avg. FD** | 0.011165 | 0.85851 | 0.6226 |
| **Occipital lobe avg. FD** | 0.0048239 | 0.43259 | 0.6666 |
| **Cingulate/insula avg. FD** | 0.010144 | 0.68106 | 0.62260 |

### Greater BSC in individuals with ASD

We observed regions of significantly greater BSC in individuals with ASD compared to controls in the bilateral superior temporal gyrus, inferior temporal gyrus, and left inferior frontal gyrus (<5% FDR, peak Cohen’s d = 0.36) (Fig. 2a), corresponding to a faster transition from cortical gray matter to white matter relative to controls. Effect sizes varied moderately between sites but were mostly positive (Fig. 2b, Supplementary Fig. S10). The analysis of the main effect of ASD was repeated with a smoothing kernel of 10mm FWHM, where the pattern of group differences is similar, and more extensive (Supplementary Fig. S11).

**Figure 2.**
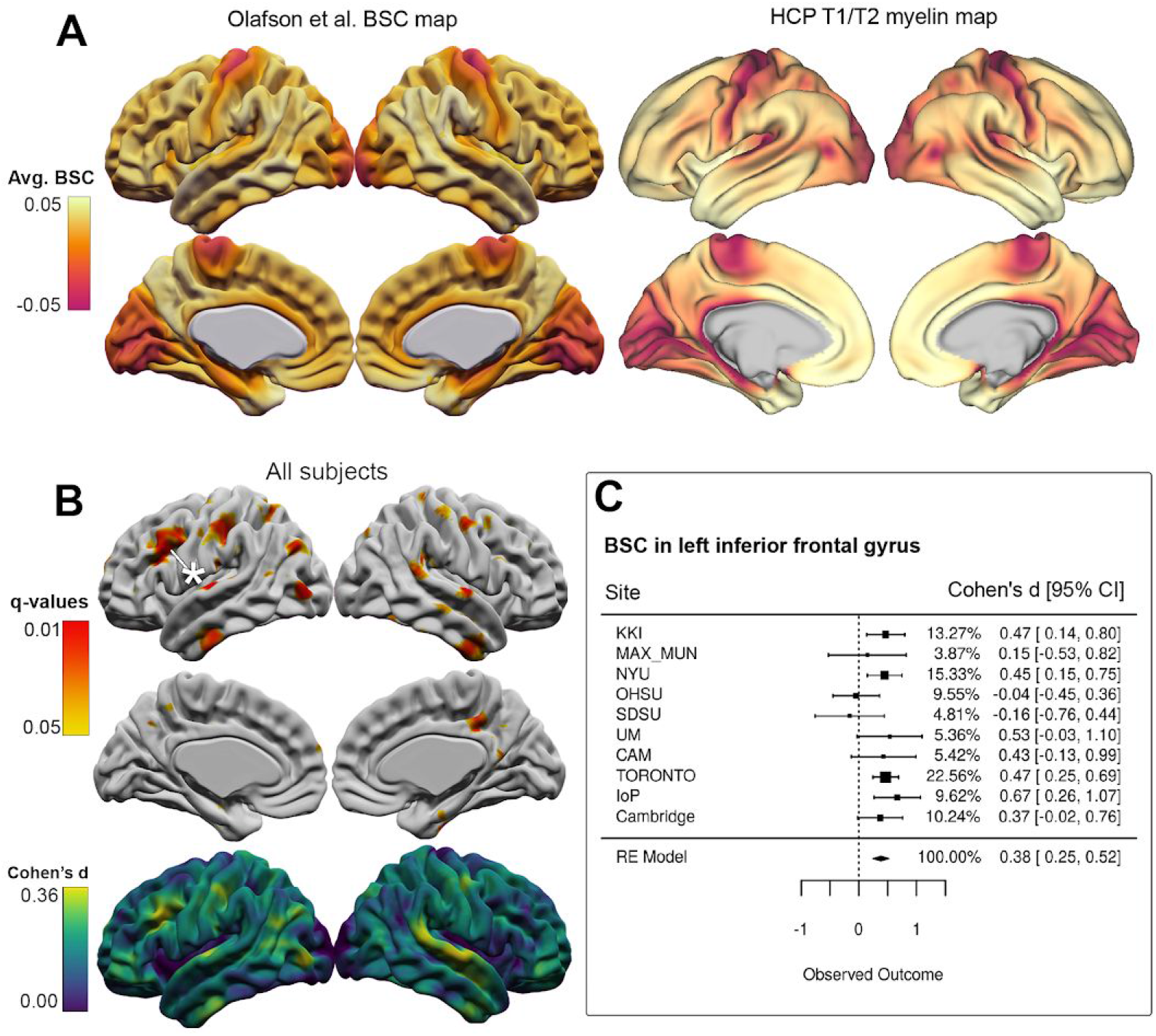
Diagnostic group comparisons of BSC. (A) Average BSC map versus average T1/T2 ratio myelin map from Glasser et al., (2011). Olafson et al. BSC map depicts the average BSC values across control subjects aged 22-35 for a single site (Sick Kids) for correspondence with the HCP dataset used by Glasser et al. (2011) to derive T1/T2 ratio maps. The top row displays lateral views (left hemisphere on the left, right hemisphere on the right), and the bottom row displays medial views (with midline vertices masked out), for each map. (B) Individuals with ASD had significantly higher BSC measures (<5% FDR, peak Cohen’s d = 0.38) in the bilateral superior temporal gyrus, inferior temporal gyrus, and left inferior frontal gyrus. (C) Forest plot displaying site-specific effect sizes at a peak vertex in the left frontal gyrus represented by an asterix in (B).

### Age-specific patterns of boundary alterations

In the age-stratified analysis, individuals with ASD above and below 18 both showed significantly greater BSC than their typically developing counterparts, though these group differences showed age-specific patterning and effect sizes, and the main effect of diagnosis was stronger in the age group over 18. Individuals with ASD under 18 had significantly greater BSC in the bilateral superior temporal gyrus and right postcentral gyrus, with a peak Cohen’s d of 0.41 (Fig. 3a). Individuals with ASD over 18 had significantly increased BSC in the bilateral precuneus and superior temporal gyrus, with a peak Cohen’s d of 0.62 (Fig. 3b). In the age-centered analysis, which examined group differences at specific ages, group differences were greatest between the ages of 12 and 20 in the right superior temporal gyrus and left inferior temporal gyrus (Fig 3c, d). Threshold effects were not responsible for the pattern of differences between the two age groups (Supplementary Fig. S12a).

**Figure 3.**
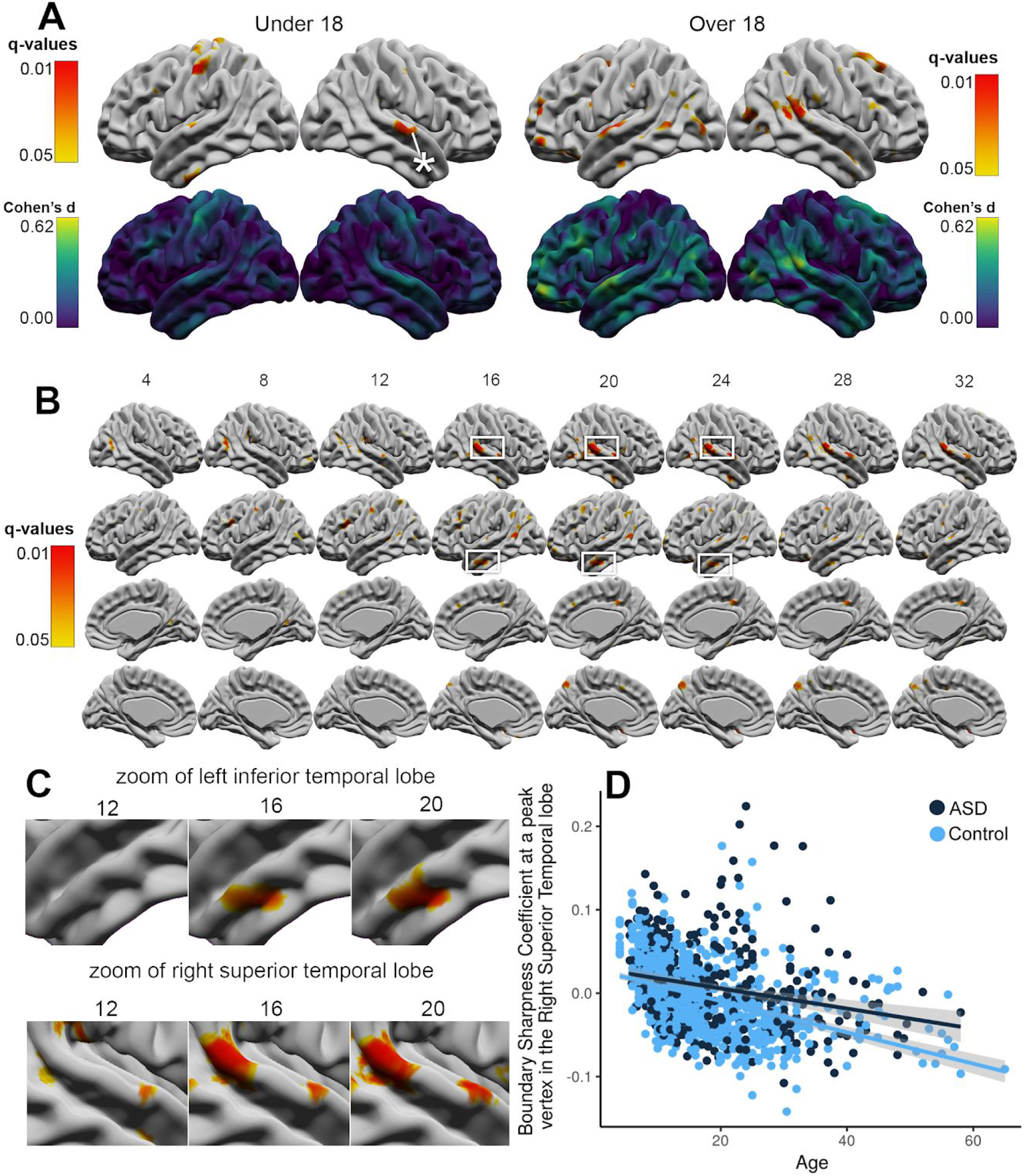
Age-stratified and age-centered analyses. Individuals under 18 with ASD had significantly higher BSC measures in the bilateral superior temporal gyrus as well as the left precentral gyrus (<5% FDR, peak Cohen’s d = 0. 41) (A). Individuals over 18 with ASD showed significantly increased BSC in the bilateral precuneus and superior temporal gyrus (<5% FDR, peak Cohen’s d = 0.62). (B) For the age-centered analysis, group differences in BSC were greatest between the ages of 12 and 20 in the right superior temporal gyrus and left inferior temporal gyrus. (D) Zoom of the left superior temporal lobe (top) and right inferior temporal lobe (bottom). (E) Plot of BSC across age in a single site (Toronto) at vertex highlighted with a white asterix in A.

### Sex-specific boundary alterations

A sex-focused analysis group differences in BSC was performed (Supplementary Methods S8). In both the male and female subgroups, individuals with ASD had significant increases in BSC, but these increases were observed in different brain regions in males and females and with larger effect sizes in the female group (Supplementary Fig. 14a, b). Females with ASD had significantly greater BSC in the bilateral superior parietal gyrus and superior temporal gyrus, with a peak Cohen’s d of 0.63 (Supplementary Fig. 14a). Males with ASD displayed significantly greater BSC in the bilateral inferior temporal gyrus and left inferior frontal lobe, with a peak Cohen’s d of 0.32 (Supplementary Fig. 14b). Threshold effects were not responsible for the pattern of differences between the sexes (Supplementary Fig. S12b).

**Figure 4.**
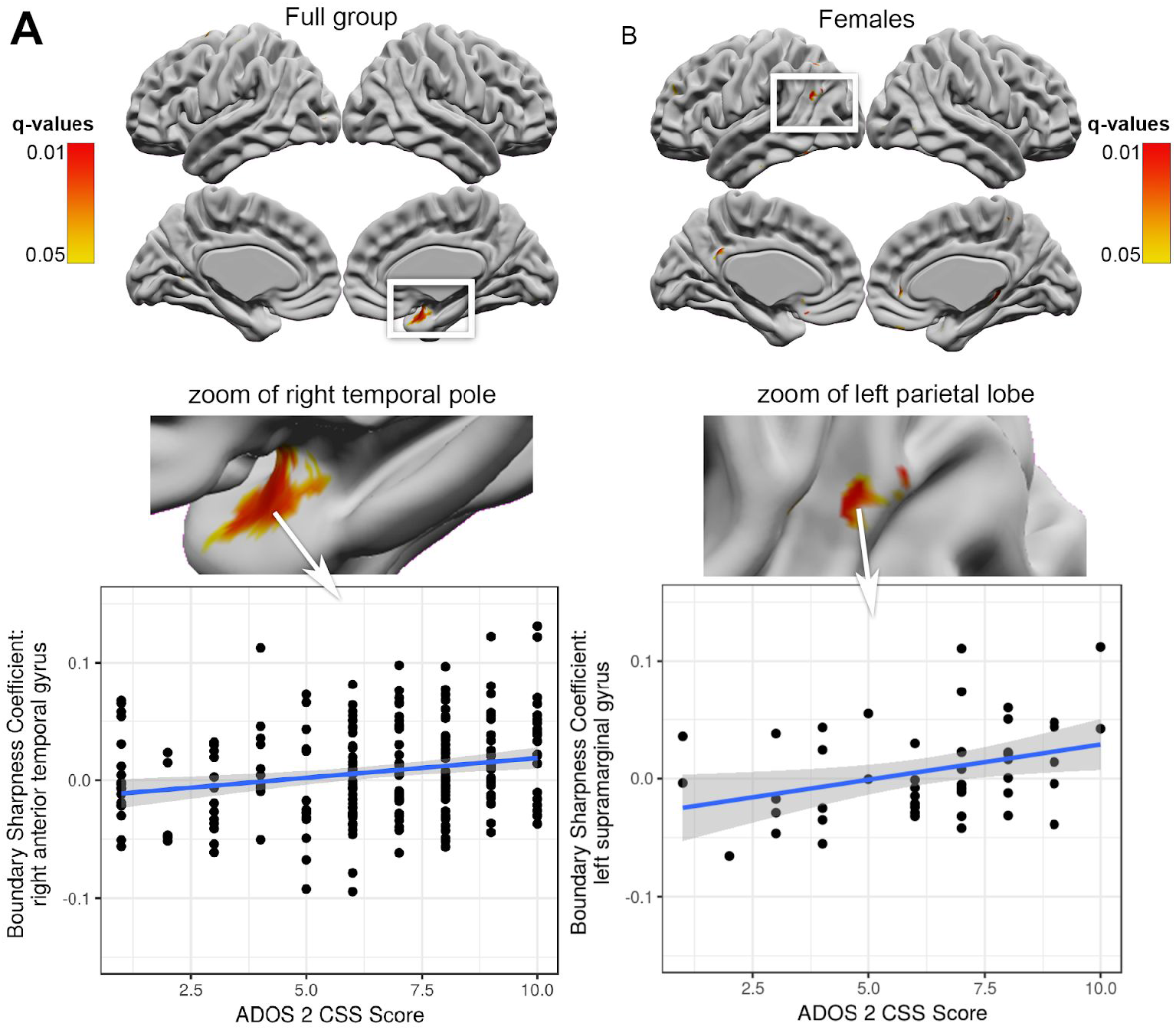
Relationship between BSC and ADOS-calibrated severity scores (CSS). Across all subjects with ASD with severity scores, ADOS-CSS was positively correlated with BSC, shown for a peak vertex in the right medial temporal gyrus (A). Correlations between ADOS-CSS and BSC were also observed in the female-only group in the left parietal lobe (B).

### Minimal correlation between BSC and ASD severity measures

We observed a significant positive correlation between BSC and ADOS-2 CSS in individuals with ASD in the right medial temporal pole (Fig. 4a) Given our findings of sex-specific regions of BSC differences in subjects with ASD, we explored the relationship between BSC and ASD severity separately in males and females. In the female group, we found a significant positive association between BSC and ADOS-2 CSS in the left parietal lobe. No correlations between severity score and BSC were observed in the male-only subset.

### Overlap between maps of diagnostic effect on cortical thickness and BSC

We observed a significant overlap between the maps of Cohen’s d effect size of diagnosis on BSC and the map of Cohen’s d effect size of diagnosis on CT (p < 0.01) (Fig. 5a,c), as well as between the maps of FDR-corrected q-values (p < 0.01) (Fig. 5b, Supplementary Fig. S13).

**Figure 5.**
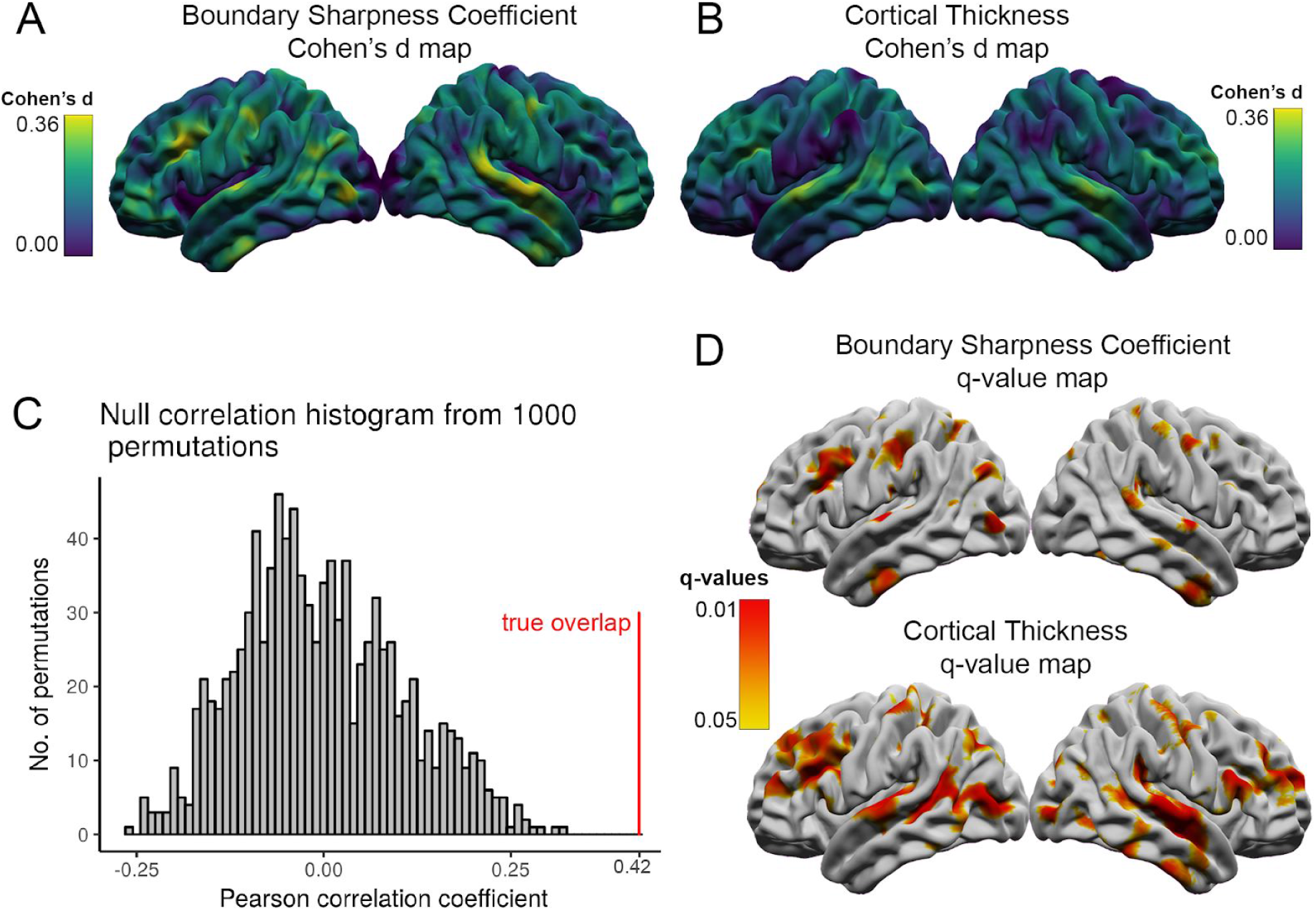
Cohen’s d effect size maps of BSC (A) and cortical thickness (B - used with permission, from Bedford et al., 2019). Spatial correspondence assessed by the Pearson correlation coefficient in a permutation-based ‘spin-test’ analysis between BSC and cortical thickness is demarcated in red (C) was significant (p= 0.00; p<<0.001 as per the software output) with 1000 null spatial permutations. FDR-thresholded q-value maps of significant increases in BSC and increases in cortical thickness in individuals with ASD (D).

## Discussion

In this study we employed a vertex-wise meta-analysis on a large multi-site dataset to investigate BSC in ASD, finding significant group-level increases in BSC in lateral frontal and temporal regions as well as sex-specific and age-specific patterns of BSC increases in individuals with ASD. As BSC is parameterized to capture cortical blurring which may be a product of differences in neural migration and intracortical myelination, there are several considerations for interpreting our observation of increased BSC in ASD. Neuronal migration defects are predicted to result in supernumerary neurons in the white matter compartment directly below the cortex (Andrews et al. 2017; Avino and Hutsler 2010; Chun and Shatz 1989). In previous neuroimaging studies examining cortical blurring at the gray/white boundary using the tissue intensity ratio, the presence of ectopic neurons is thought to be captured by an intensity differential that is lower in individuals with ASD (Andrews et al. 2017). Thus, our finding of a greater intensity difference (as indexed by greater BSC) between cortex and white matter is unlikely to reflect differences in neuronal migration.

Since T1w intensity is thought to reflect underlying myeloarchitecture more so than cytoarchitecture (Eickhoff et al. 2005), BSC may also reflect intracortical myelination: a reduction of myelin content (and presumably, of T1w signal) in the lower layers of the cortex would appear in T1w MRI as a sharper transition in intensity moving from gray to white matter. This assumption is supported by the general agreement between the group-average BSC map (Supplementary Fig. S8) and the intracortical myelin map derived using the T1w/T2w ratio by Glasser and Van Essen (Glasser and Van Essen 2011) and is further supported by evidence from studies suggesting altered intracortical myelination in ASD, including genetic (Richetto et al. 2017) and molecular (Lee et al. 2019; Canali et al. 2018) studies, preclinical mouse models (Graciarena et al. 2018; Shen et al. 2018), and postmortem histology (Zikopoulos and Barbas 2010). Furthermore, age-related effects of BSC in healthy controls (Supplementary Figure S3) recapitulate normative developmental patterns of intracortical myelination. The results suggest that BSC follows an inverted-U shape trajectory across most of the cortex, which is consistent with studies of intracortical myelin, which describe cortical development over the life span as occurring in phases: an early maturation phase that lasts into adulthood, followed by a stable period, and then a decline of intracortical myelin (Rowley et al. 2017; Grydeland et al. 2019). Here, across most of the cortex, BSC trajectories follow a similar pattern - an early decrease in BSC (increase in myelin) during adolescence, and then a plateau, and then a moderate increase in BSC (decrease in myelin).

Mounting evidence from diffusion imaging and resting state fMRI studies support the characterization of ASD as a connectopathy, or a brain network disorder. Since our proposed measure is parameterized to capture myelination in the lower layers of the cortex (a region through which fibers that connect distant brain regions pass), the higher BSC observed in ASD is supported by studies that find reduced long-range cortical connectivity (Kikuchi et al. 2015; O’Reilly, Lewis, and Elsabbagh 2017) and thalamo-cortical connectivity (Tomasi and Volkow 2019; Nair et al. 2013) in ASD. As such, the contribution of myelination is increasingly relevant to understanding the neurodevelopmental mechanisms of ASD.

Two neuroimaging studies to date investigated boundary microstructure in ASD using a different metric called the tissue intensity ratio, reporting lower tissue intensity ratio in ASD which may indicate greater intracortical myelination or the presence of supernumerary neurons in the superficial white matter compartment directly beneath the gray/white boundary (Andrews et al. 2017; Mann et al. 2018; Norbom et al. 2019). This discrepancy to the present findings could be due, in part, to differences in preprocessing and analysis methods (particularly the use of the tissue intensity ratio). The tissue intensity ratio suffers from the ambiguity arising from the blurring around the cortical boundary which, in turn, alters placement of the gray/white boundary that serves as a reference point for the component gray and white measurements. Additionally, the present study uses a sample size that is considerably larger than previous studies and is more suitably powered to detect group differences in high-variability populations. Examining the directionality in each of the forest plots reveals how different studies may, indeed, yield different findings and how they may be dependent on age range and demographic composition. The data in the present study were rigorously filtered for motion and image processing artefacts (Bedford et al. 2019) whereas quality control procedures were not described in detail in previous studies, making it difficult to assess the effect of motion or inaccurate segmentation on reported results. It is possible that measures of cortical blurring using the tissue intensity ratio in previous ASD studies have been obscured by motion-induced blurring of the gray-white matter boundary (A. Alexander-Bloch et al. 2016; Reuter et al. 2015), especially considering individuals with ASD are more likely to move during the scan than controls (Pardoe, Kucharsky Hiess, and Kuzniecky 2016; Bedford et al. 2019). It is also possible that factors such as age, sex, FIQ, and symptom severity are influencing case-control differences in previous studies of cortical blurring in autism (Lombardo, Lai, and Baron-Cohen 2019). Therefore, we attempted to determine the degree to which diagnostic differences in BSC are modulated by these factors.

The male bias in ASD prevalence (3:1 males:females diagnosed (Baxter et al. 2015) as well as sex differences in behaviour and key ASD-related phenotypes such as restrictive and repetitive behaviours (Knutsen et al. 2019; Mandy et al. 2012) have spurred the investigation of neuroanatomical sex differences in ASD (M. C. Lai et al. 2017). Given their scarcity, females with ASD are a difficult population to recruit, and, as such, our understanding of the modulating effect of sex on neuroanatomy in individuals with ASD is relatively rudimentary. However, this knowledge gap is beginning to narrow as a result of efforts to curate large-scale publicly-available neuroimaging datasets (A. Di Martino et al. 2014) and an increased awareness of the importance of female representation in ASD studies (M.-C. Lai et al. 2015). By combining publicly-available datasets with several multicentre consortium datasets, we were able to incorporate data from 117 females with ASD in this study. We observed sex-specific patterns of BSC, where females with ASD displayed higher BSC in the superior temporal and parietal lobes, whereas males with ASD displayed a greater BSC in the inferior temporal and frontal lobes. Sex-specific patterns of cortical neuroanatomy in autism have recently been reported, in both gross volumetric measures (Bedford et al. 2019; Retico et al. 2016) and altered connectivity (Irimia et al. 2017; Zeestraten et al. 2017). The maximum effect size of diagnosis in the female sample was almost twice as high as the effect size of diagnosis observed in the male sample. BSC also correlated with ADOS-CSS in the female-only sample. Taken together, these findings support a differential neuroanatomical presentation of autism in males and females.

Age is another factor that is known to modulate diagnostic group differences. Though most brain differences in ASD have been investigated in a child or adolescent population, there is evidence that differences in brain anatomy related to ASD are present even in adulthood (Lazar et al. 2014; Ecker et al. 2013, 2012). Intracortical myelination has shown to be ongoing even past adolescence, with accelerated myelination until ∼30 years of age, followed by a period of stability, and then a decrease in myelination from the late 50s (Tullo et al. 2019; Grydeland et al. 2013). Though our use of cross-sectional data limits our interpretation of the age-centered results as being reflective of developmental processes, the peak diagnostic group differences observed in adolescence and adulthood may be the result of reduced or protracted myelination in ASD relative to the rate of myelination in typical development, as observed in postmortem histology (Zikopoulos and Barbas 2010) and mouse models of ASD (Graciarena et al. 2018; Ellegood et al. 2015).

Intracortical myelination has been shown to drive MRI-based measures of cortical thickness in the visual cortex across development (Natu et al. 2018), potentially by pushing the gray-white matter boundary deeper into the cortex. In our analysis, we found a significant spatial overlap between maps of the effect size of diagnosis on BSC and the effect size of diagnosis on cortical thickness (Bedford et al. 2019). These results suggest that the neuro-phenotype of increased cortical thickness in ASD, as observed in (Bedford et al. 2019) and many other studies, may be partially driven by lower levels of intracortical myelination in ASD relative to controls. Moving forward, a longitudinal design would allow us to determine in the same individual, the evolution of the pattern of BSC across development, and would allow for relating patterns of BSC over time with respect to behaviours and autism symptomatology. Additionally, since age may substantially moderate the pattern of sex differences, a longitudinal framework would allow for a more precise investigation into this time-sensitive relationship.

However, while cortical thickness differences observed in our previous work (Bedford et al. 2019) correlate with autism severity scores in many regions of the brain, we found minimal correlations between BSC and ASD severity. This disparity suggests that intracortical myelination may inflate MRI-based measures of cortical thickness in ASD for several areas of the brain, but the cortical overgrowth phenotype itself is driven by separate biological processes (such as reduced synaptic pruning) that are more clinically relevant to ASD severity. On the other hand, deficits in intracortical myelination may be a neurobiological hallmark of the disorder that does not alter ASD behavioural severity beyond a certain point, and thus does not correlate with the ADOS-CSS score.

There are several limitations to this study. First, although our BSC maps correspond well with established patterns of intracortical myelination, the T1w intensity also reflects other biological properties including water content, iron, and dendrite density, and therefore may reflect more than the degree of myelination in the cortex (Stüber et al. 2014). Second, abnormalities in the superficial white matter may also drive some of the changes to BSC. Though a vertex-wise analysis of the intracortical or superficial white matter intensities independently is precluded by between-scanner intensity profile differences and by the distorting effect of inhomogeneity gradients across images, it would be possible to evaluate differences in normalized intracortical intensities that have been scaled to the intensity of a predetermined sample of white matter within each subjects’ brain. Finally, since the distance sampled in the white matter is based on a percentage of the cortical thickness, the depth at which white matter is measured varies between regions with different cortical thicknesses.

Given these findings of increased BSC in individuals with ASD, which may reflect the degree of myelination in the lower layers of the cortex, it will be pertinent to investigate how subcortical volume changes in ASD relate to BSC in cortical sensory regions with high thalamic input (Schuetze et al. 2016; Uddin 2015). Considering the importance of intracortical myelination on the fidelity of neural connections and the maintenance of networks, these results may help us better understand the cognitive and behavioural atypicalities seen in ASD. Finally, the significant spatial correspondence observed between maps of cortical thickness and of BSC calls for a reconsideration of what biological phenomena may underlie the MRI-derived measures of cortical thickness increases in autism.

## Supplementary Information

### Supplementary Methods

#### S1. Sample details

##### S1.1 Data processing

All data used in this study was preprocessed by SB for analysis in Bedford et al., (2019), and sample characteristics (including image processing, and which subjects were excluded due to poor motion or CIVET quality) are equivalent between studies, with the exception of the NIMH site, which was not included here. See Bedford et al. (2019) for imaging parameters and preprocessing considerations for each site. The data used in this study was obtained from scans whose CIVET/1.1.12 segmentations quality had already been assessed. As such, a newer version CIVET was not re-run on this data.

##### S1.2 Quality Control

###### S1.2.1 Motion QC

In-scanner movements cause motion artefacts in MR images, which can influence certain morphological measurements like cortical thickness and cortical volume (A. Alexander-Bloch et al. 2016; Reuter et al. 2015). As such, performing quality control (QC) assessments for motion artefacts is essential to eliminate statistical bias in disorders associated with movement abnormalities, such as ASD. All data in this report was rated for motion by three independent raters (SB, MMC, and ST). Data from subjects with excessive motion scores were excluded from analyses (Bedford et al. 2019). Specifically, raw images were rated on a 4-point scale based on the presence of ringing or blurring, and any motion or acquisition-related artefacts in the image (1 = no motion; 2 = slight motion; 3 = obvious motion; 4 = very bad or extensive motion). After training on a QC manual, inter-rater reliability was assessed after the first 100 images were scored, and discrepancies were addressed through consensus discussion.

###### S1.2.2 CIVET QC

In addition to motion QC, data from subjects whose CIVET QC scores were below a certain threshold were excluded (Bedford et al. 2019). Specifically, white and gray matter surface accuracy was rated by 2 independent raters using a CIVET-specific QC manual. As there were only 3 levels of quality control in this case, a discrepancy was considered any two scores that were not in perfect agreement between the two raters. Outputs were given a score of 0 (fail), 0.5 (warn), or 1 (pass). A score of 1 was given for outputs with accurate classification and boundaries, and no visible errors. A 0.5 was given for an overall good segmentation with a small error confined to a small region of cortex, such as under-segmentation or minor classification errors. A score of zero was given for larger errors or multiple small errors. In total, after CIVET and motion QC, there remained 1056 subjects: 427 with ASD (310 male/117 female) and 629 controls (323 male/306 female).

###### S1.2.3 Demographics of patients excluded due to motion and CIVET QC

As determined by Bedford et al. (2019), participants excluded due to motion and CIVET failures were significantly younger (p=2.2e-16), had lower FIQ (p=4.2e-08), and included higher numbers of males (p=4.8e-11), and individuals with ASD (p=3.3e-08). ASD individuals that failed QC had significantly higher levels of autistic symptoms based on the ADOS-G total scores (p=2.5e-06), but there were no significant differences in ADOS-2 CSS (p=0.20). Excluded participants with ASD also had higher levels of restricted, repetitive behaviours, as measured by the ADOS-G (p=0.0001) and the ADI-R: p=0.0008).

#### S2. Surface Generation

##### S2.1 New gray matter surfaces

A gray matter surface at 25% of the distance from the gray/white boundary surface to the pial surface was generated by averaging the vertices of the gray/white boundary surfaces (0%) and mid-surfaces (50%), which were created in standard space by CIVET/1.1.12, using the function *average_surfaces*. For each subject, the 12.5% gray matter surfaces were generated by combining the 0% surface with the 25% surface, 6.25% surfaces by combining the 0% surface with the 12.5% surface, and 18.75% surfaces by combining the 12.5% surfaces with the 25% surfaces.

##### S2.2 New white matter surfaces

Each new white matter surface was generated by first creating vectors between each vertex on the gray/white boundary surface and the corresponding vertex on the gray matter surfaces (repeated separately for each gray matter surface generated (S2.1), with the exception of the 50% or mid-surface). The vectors were then inverted in the x, y, and z dimensions and added to the coordinates of the graywhite boundary surface vertices, such that white matter surfaces were created at the same distance from the gray/white boundary as the gray matter surfaces, but in the opposite direction (toward the superficial white matter instead of toward the pial surface).

#### S3. Model fit

A sigmoid curve (Equation 1) was fit to the 10 sampled intensities by finding the optimal least squares solution over parameters (a, k, c, d) between a[min(yvalues), 2000], k[0, 100], c[0,1], and d[-50,50], using the nls() function and “port” algorithm in R (Figure S3). A base sigmoid function (Weisstein n.d.) was reparameterized to constrain the shape of the curve to be increasing from the pial surface to the superficial white matter. In a single-subject analysis, convergence was achieved for ∼98% of vertices that were regularly distributed across the brain (Figure S4).

#### S4. Tissue Intensity Ratio

The tissue intensity ratio was calculated by dividing the gray matter intensity measured at 25% of the cortical thickness by the white matter intensity measured at an equal distance (25% of the cortical thickness) in the direction of the white matter. Values were resampled to a common mesh and smoothed, except for the single-subject analysis (Supplementary Fig 1), in which values were not resampled before smoothing so as to not introduce correlations with the curvature that are used to guide surface-based registration.

#### S5. Statistical models

For the case-control comparisons, the following model was used, with i representing each vertex across the brain:

**Case-control analysis**: BSC_i_ ∼ β_0_Intercept + β_1_Age + β_2_Sex + β_3_Diagnosis + β_4_FIQ +ε

**Sex-stratified analysis**: BSC_i_ ∼ β_0_Intercept + β_1_Age + β_2_Diagnosis + β_3_FIQ + ε

**Age-centered analysis:** BSC_i_ ∼ β_0_Intercept + β_1_(Age-*a*)*Diagnosis +β_2_ Sex + β_3_FIQ +ε

Where *a* is the age at which group differences are assessed.

**Age-stratified analysis:** BSC_i_ ∼ β_0_Intercept + β_1_Age +β_2_ Sex + β_3_Diagnosis + β_4_FIQ+ ε

**Model assessment:** To test the importance of sex, age and FIQ, an Aikaike Information Criteria (AIC) was used. The AIC is a measure that assesses the relative quality of statistical models, where a model with the lowest AIC is considered the best fit for the data.

Sex:

BSC_i_ = β_0_ + β_1_Diagnosis + β_2_Sex + β_3_(Diagnosis*Sex) + β_4_Age + β_5_FIQ + ε_i_

BSC_i_ = β_0_ + β_1_Diagnosis + β_2_Sex + β_4_Age + β_5_FIQ + ε_i_

BSC_i_ = β_0_ + β_1_Diagnosis + β_2_Age + β_3_FIQ + ε_i_

Age:

BSC_i_ = β_0_ + β_1_Diagnosis + β_2_Sex + β_3_Age + β_4_(Diagnosis*Age) + β_5_FIQ + ε_i_

BSC_i_ = β_0_ + β_1_Diagnosis + β_2_Sex + β_3_Age + β_5_FIQ + ε_i_

BSC_i_ = β_0_ + β_1_Diagnosis + β_2_Sex + β_5_FIQ + ε_i_

Age-squared:

BSC_i_ = β_0_ + β_1_Diagnosis + β_2_Sex + β_3_Age + β_4_Age^2^ + β_5_(Diagnosis*Age^2^) + β_6_FIQ + ε_i_

BSC_i_ = β_0_ + β_1_Diagnosis + β_2_Sex + β_5_Age + β_4_Age^2^ + β_4_FIQ + ε_i_

BSC_i_ = β_0_ + β_1_Diagnosis + β_2_Sex +β_5_Age + β_4_FIQ + ε_i_

FIQ:

BSC_i_ = β_0_ + β_1_Diagnosis + β_2_Sex + β_3_Age + β_4_FIQ + β_4_(Diagnosis*FIQ) + ε_i_

BSC_i_ = β_0_ + β_1_Diagnosis + β_2_Sex + β_3_Age + ε_i_

#### S6. Regression of age, age-squared, and sex on BSC in controls

A vertexwise regression of age, age squared, and sex on BSC across all healthy individuals in the Sick Kids (Toronto) cohort was performed, followed by false discovery rate correction.

#### S7. In-scanner motion and BSC

In-scanner motion was approximated by average Framewise Displacement (FD) (Power et al. 2012). Rotational displacements were converted from degrees to millimeters by calculating displacement on the surface of a sphere of radius 50 mm. The following regression model was performed separately for each lobe (left and right hemisphere averaged) and for the whole brain:

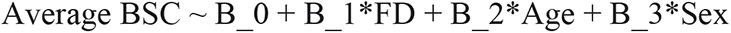

#### S8. Sex-focused analysis

Sex-specific patterns of BSC were assessed by stratifying subjects by sex and performing the same case-control meta-analysis in males and females separately, including age and FIQ as covarites.

## Supplementary Figures

**Supplementary Figure S1.**
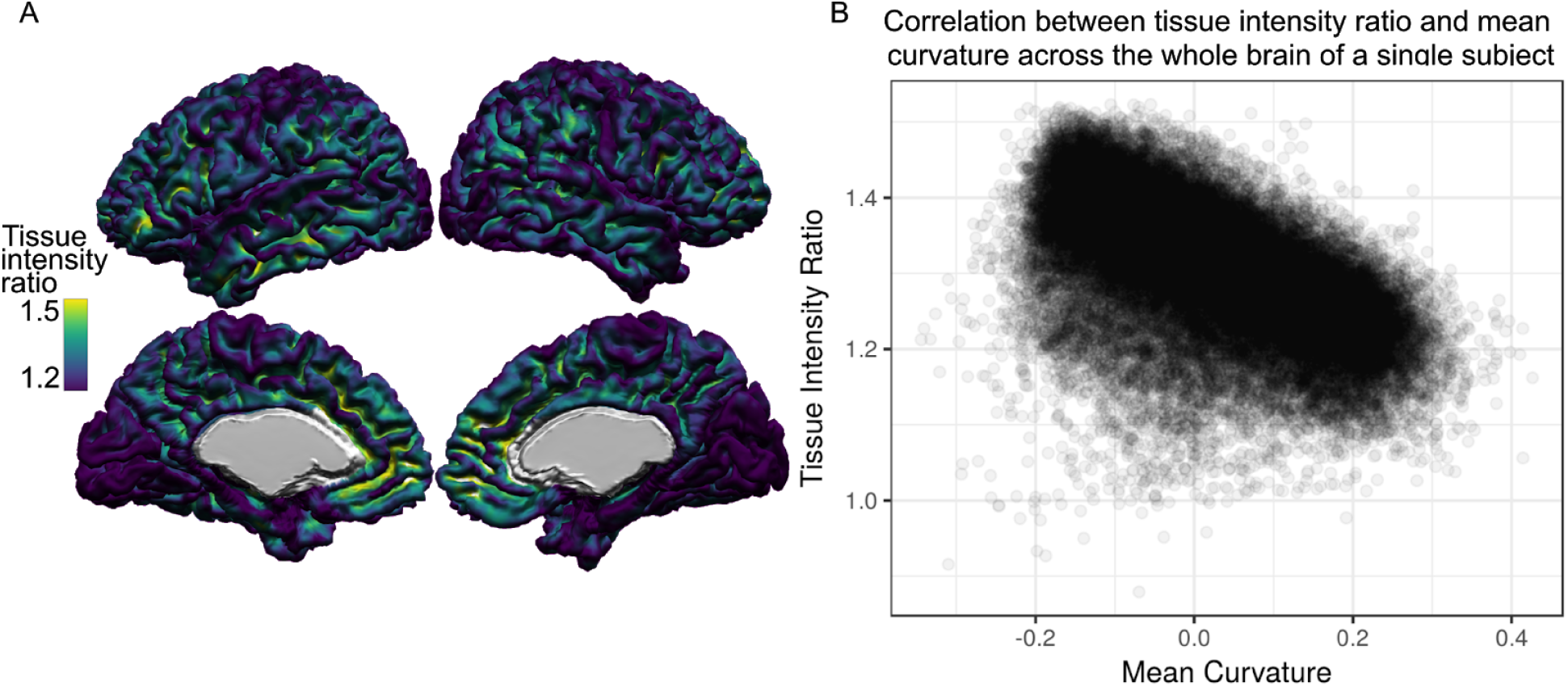
Left: tissue intensity ratio values displayed on a single subject’s mid-cortical surface (i.e. the surface at 50% of the distance between the pial and white matter surface). Right: vertex-wise correlation between the tissue intensity ratio calculated as the ratio of white matter to gray matter intensities and mean curvature. The tissue intensity ratio is significantly negatively correlated with mean curvature for all subjects (<5% FDR, data not shown) in a single exploratory site (Sick Kids; 88 ASD/314 CTL), meaning that the tissue intensity ratio is higher in convex regions, such as sulci, and lower in concave regions, like gyri. Vertex-wise correlations were assessed in the native space before resampling to a common surface mesh.

**Supplementary Figure S2.**
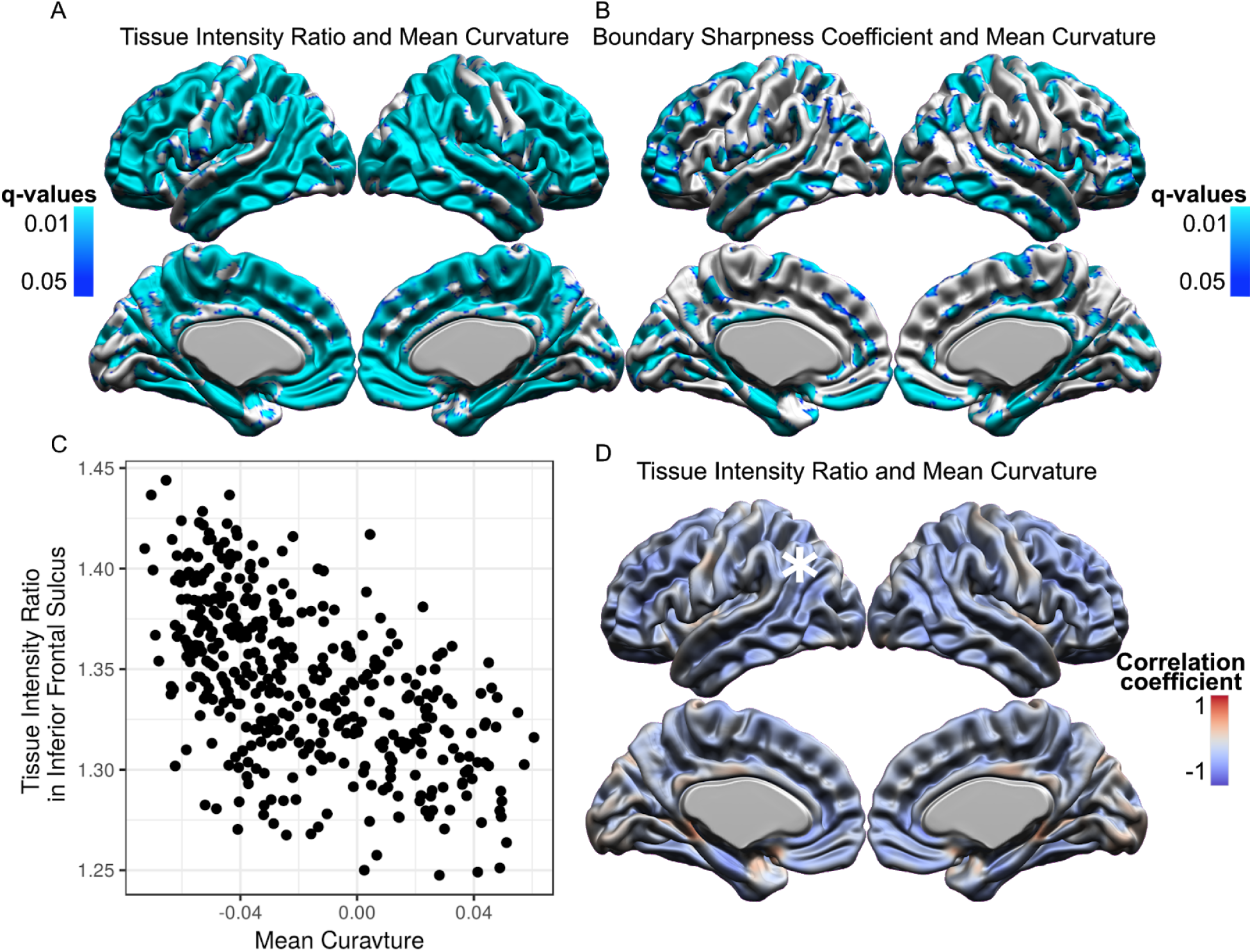
Regions of significant (<5% FDR) cross-subject correlation of the tissue intensity ratio with mean curvature (A) and of BSC with mean curvature (B) at each vertex in a single exploratory site (Sick Kids; 88 ASD/314 CTL). Relationship between mean curvature and tissue intensity ratio across subjects at a vertex in the right inferior frontal sulcus, Pearson’s correlation coefficient = -0.56 (C). Map of the correlation coefficient between the tissue intensity ratio and mean curvature (D); white asterisk on the left lateral surface represents the vertex shown in (C).

**Supplementary Figure S3.**
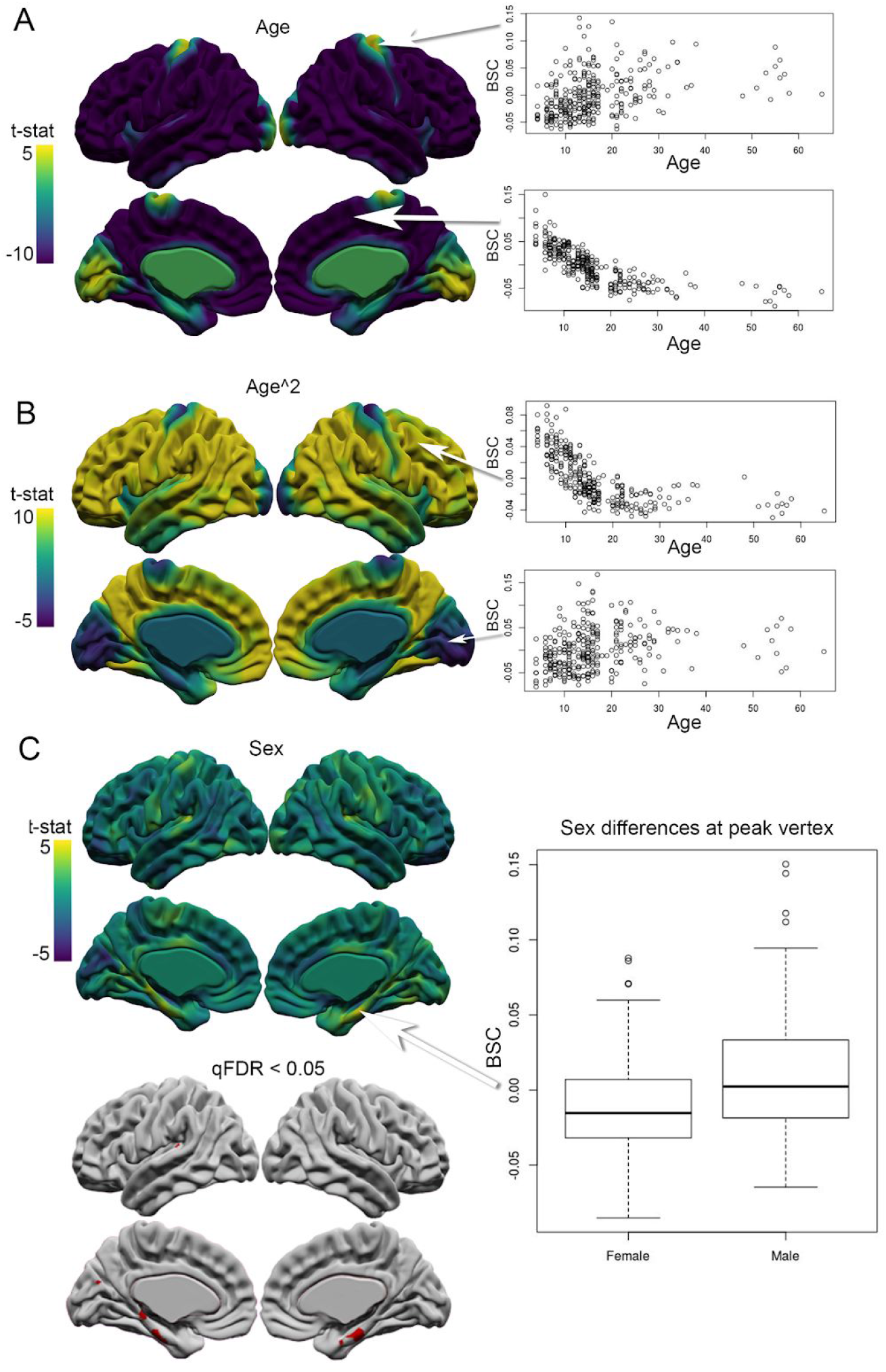
Normative effects of age, age squared, and sex, on BSC in a single cohort of healthy individuals. (A) Age is significantly related to BSC across the brain, with a positive relationship observed in primary sensory areas, and a negative relationship in all other areas. (B) Age squared also displays a significant relationship with BSC across the brain, where inverted U-shaped trajectories of age and BSC are fit to primary sensory areas, and all other cortex is better fit by a U-shaped trajectory. For age and age-squared, the subcortical midline vertices are not included. (C) Sex is significantly associated with BSC in areas of the medial temporal cortices, where males have significantly greater BSC than females; regions in which BSC is significantly modulated by sex at an FDR threshold of q < 0.05 are displayed in red.

**Supplementary Figure S4.**
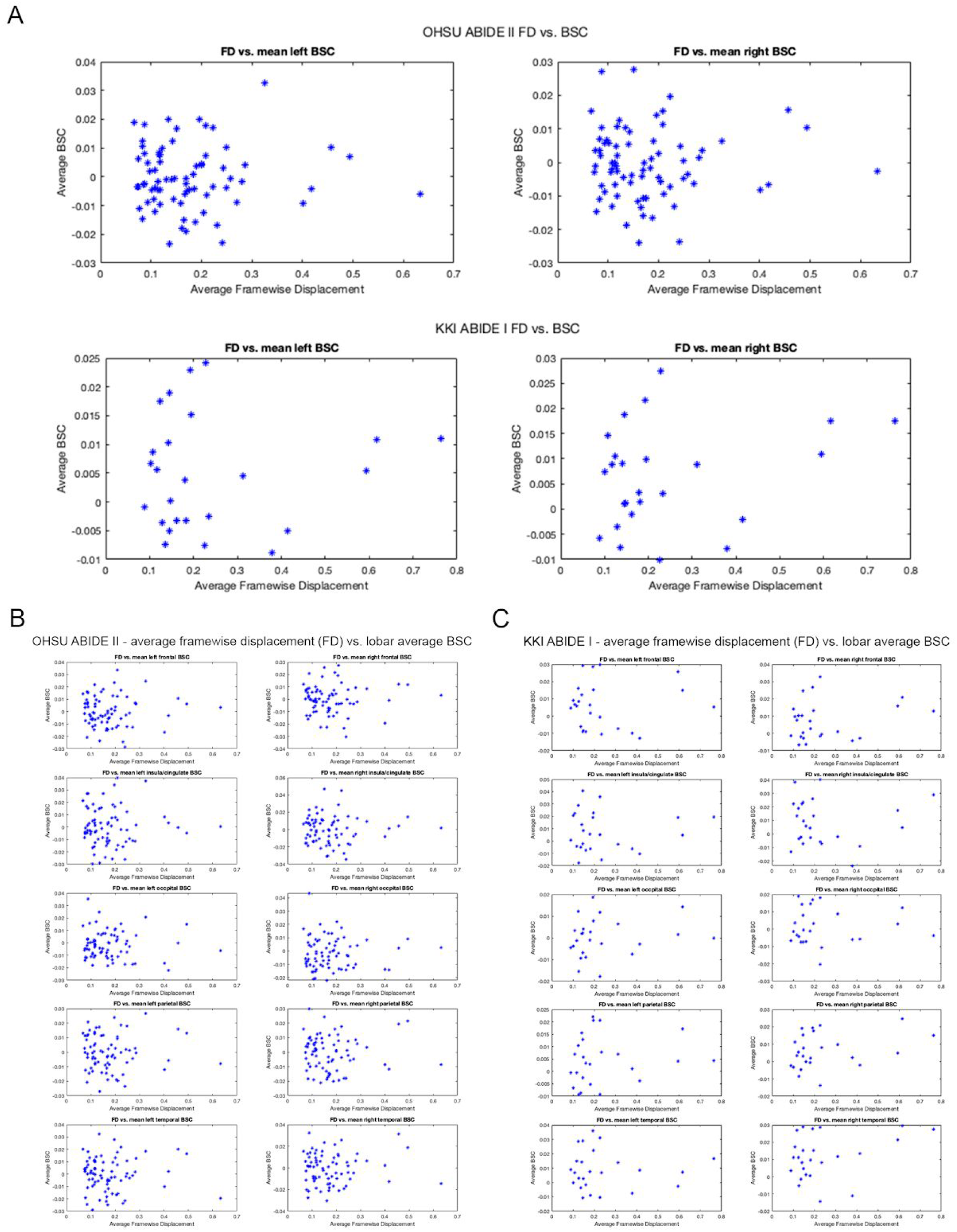
Relationship between average Framewise Displacement and (A) whole-brain average BSC and lobar average BSC in (B) ABIDE II OHSU and (C) ABIDE I KKI (each point is one subject) containing healthy controls and individuals with ASD between ages 7 and 11. None of the relationships are significant when controlling for age and sex.

**Supplementary Figure S5.**
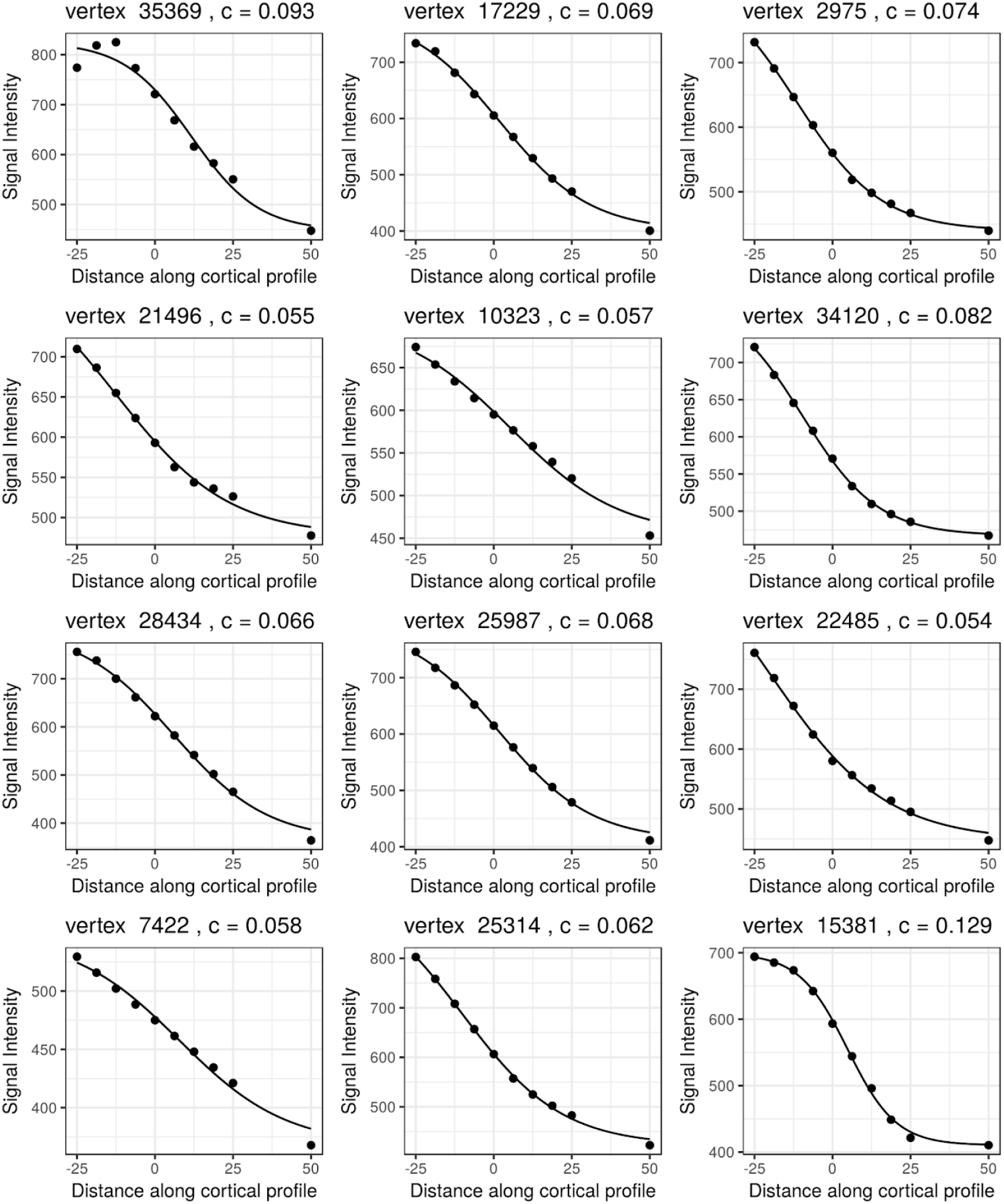
Image intensity samples (black dots) and estimated sigmoid curve (black line) from 12 randomly chosen vertices in a single subject. The x-axis displays distance along the cortical profile, starting at the outermost surface in the superficial white matter (denoted as -25%) and ending at the gray matter midsurface (50%). BSC values prior to log-transformation are displayed in the title of each plot.

**Supplementary Figure S6.**
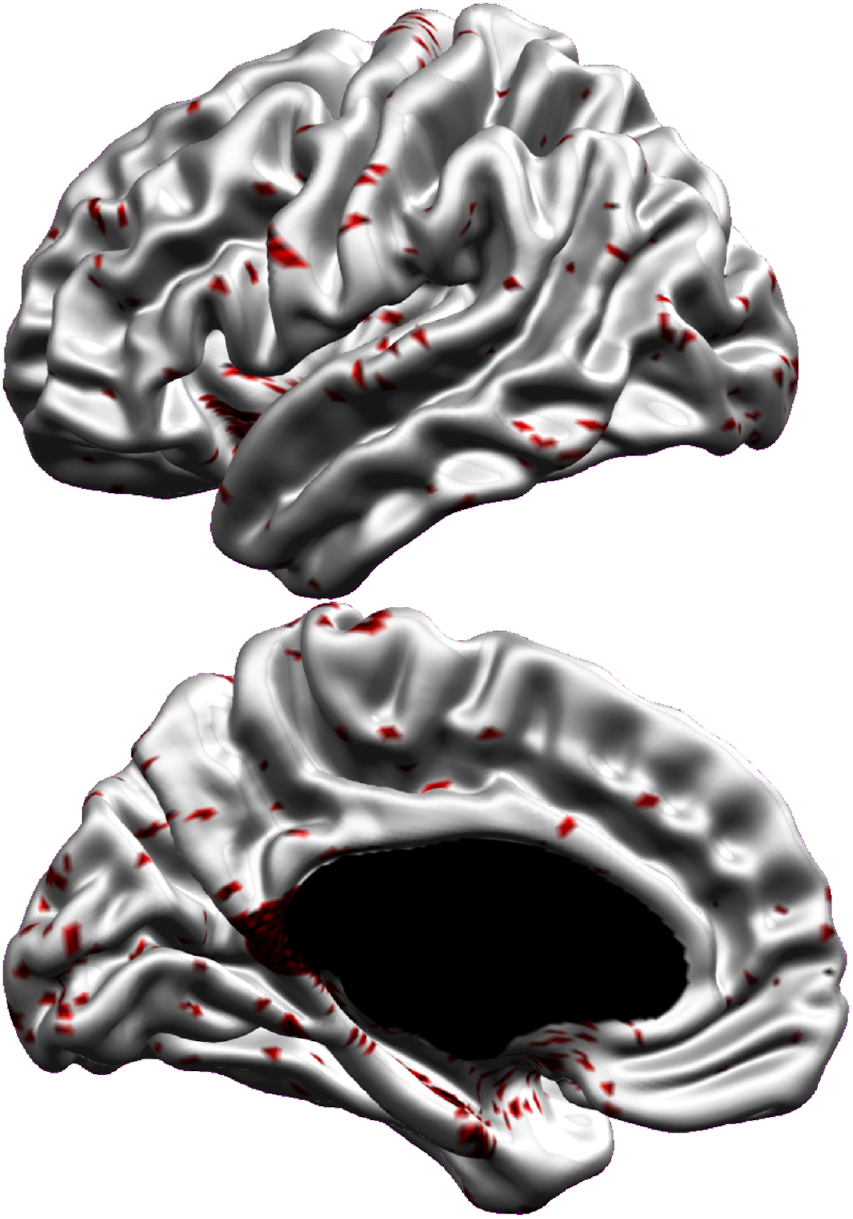
Distribution of convergence failures (i.e. where a sigmoid curve could not be adequately fit) for a single subject, shown in red. The subcortical surface is masked in black.

**Supplementary Figure S7.**
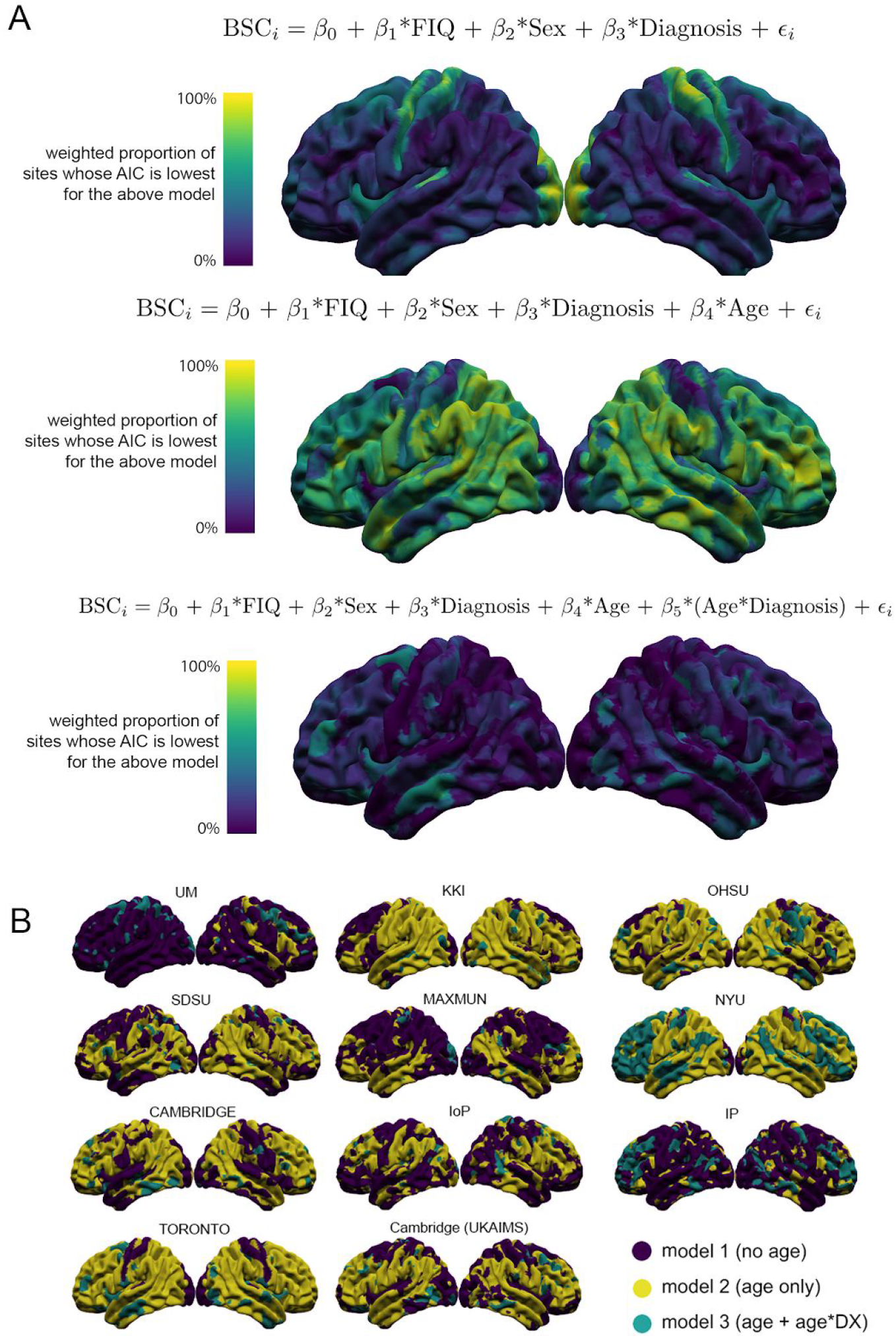
Akaike information criterion (AIC) analysis reflecting the importance of age on BSC. (A) Color maps represent the percentage of sites (calculated as a weighted average, based on site size after QC) for which BSC best explained by either the model with diagnosis only (top row), diagnosis and age (middle) and diagnosis, and, and diagnosis*age (bottom). (B) For each site, the spatial distribution of which model was best fit at each vertex.

**Supplementary Figure S8.**
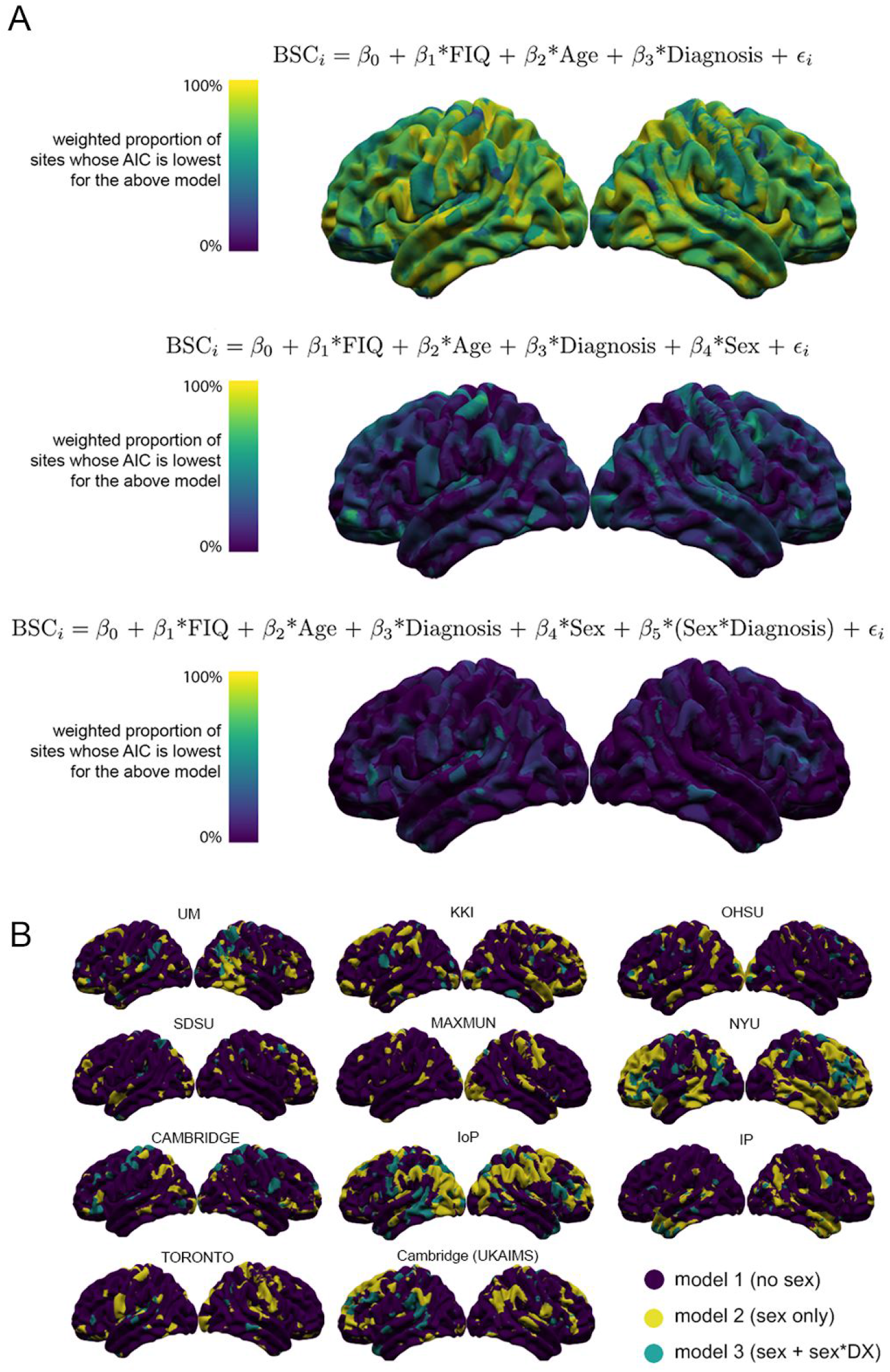
Akaike information criterion (AIC) analysis reflecting the importance of sex on BSC. (A) Color maps represent the percentage of sites (calculated as a weighted average, based on site size after QC) for which BSC best explained by either the model with diagnosis only (top row), diagnosis and sex (middle) and diagnosis, and, and diagnosis*sex(bottom). (B) For each site, the spatial distribution of which model was best fit at each vertex.

**Supplementary Figure S9.**
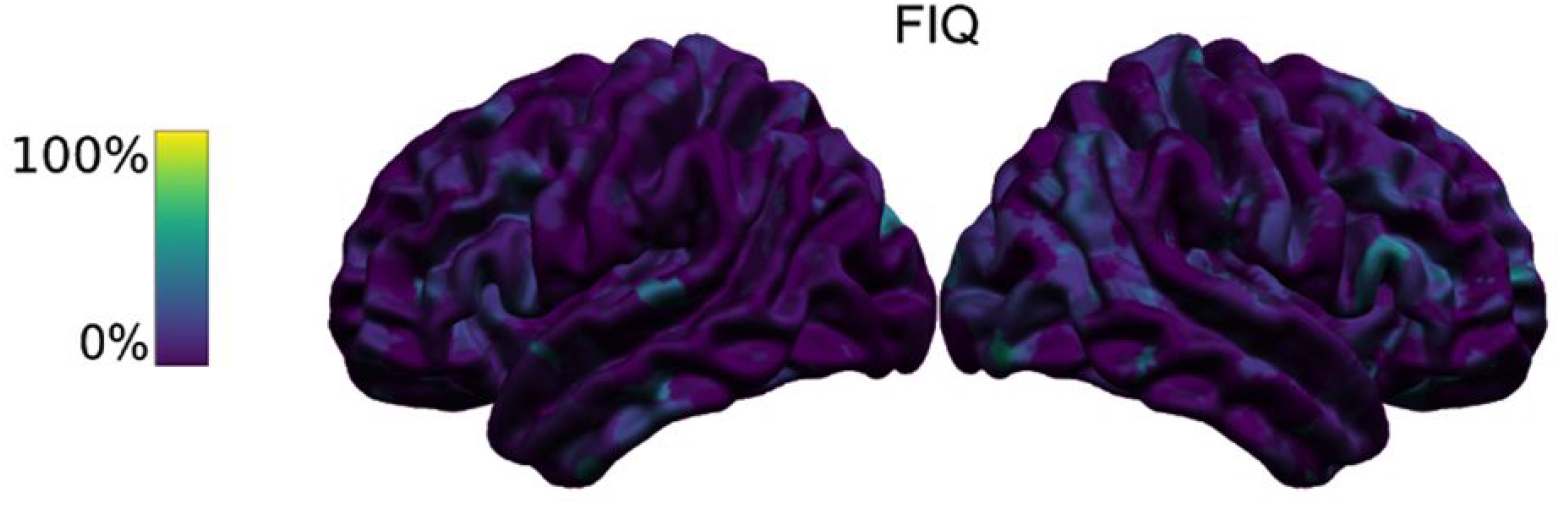
Akaike information criterion (AIC) analysis reflecting the importance of FIQ on BSC. Color map represents the percentage of sites (calculated as a weighted average, based on site size after QC) for which BSC best explained by the model with diagnosis, FIQ, and diagnosis*FIQ compared to the model with just diagnosis.

**Supplementary Figure S10.**
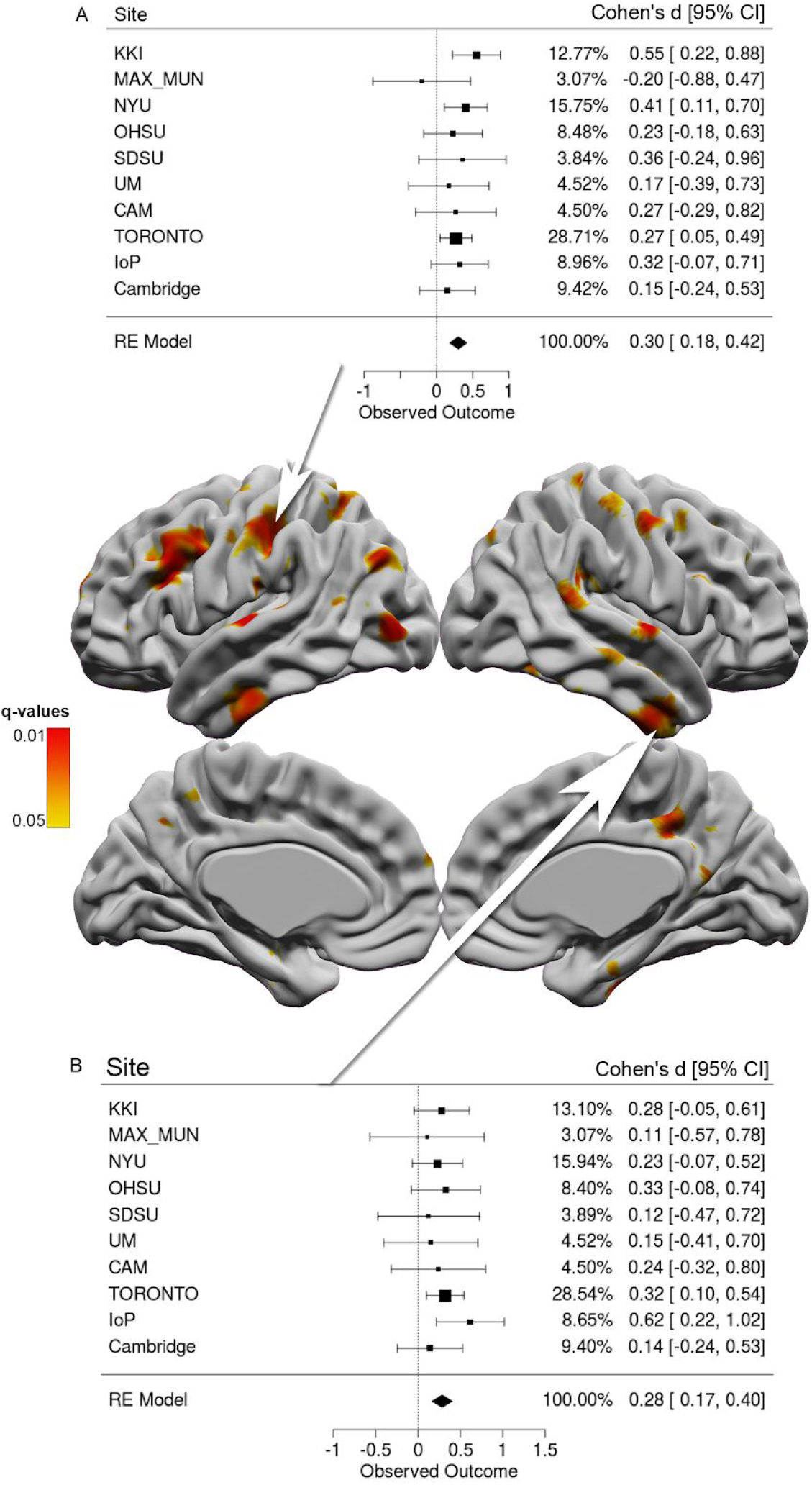
Significant vertex-wise group differences (ASD vs. control) in BSC across all individuals at 5% FDR. Forest plots show effect sizes per imaging site at two regions of interest highlighted with a red arrow: the left posterior postcentral gyrus (A) and right inferior temporal gyrus (B). Top row displays lateral views (left hemisphere on the left, right hemisphere on the right), and the bottom row displays medial views (left hemisphere on the left, right hemisphere on the right - subcortical midline structures masked out).

**Supplementary Figure S11.**
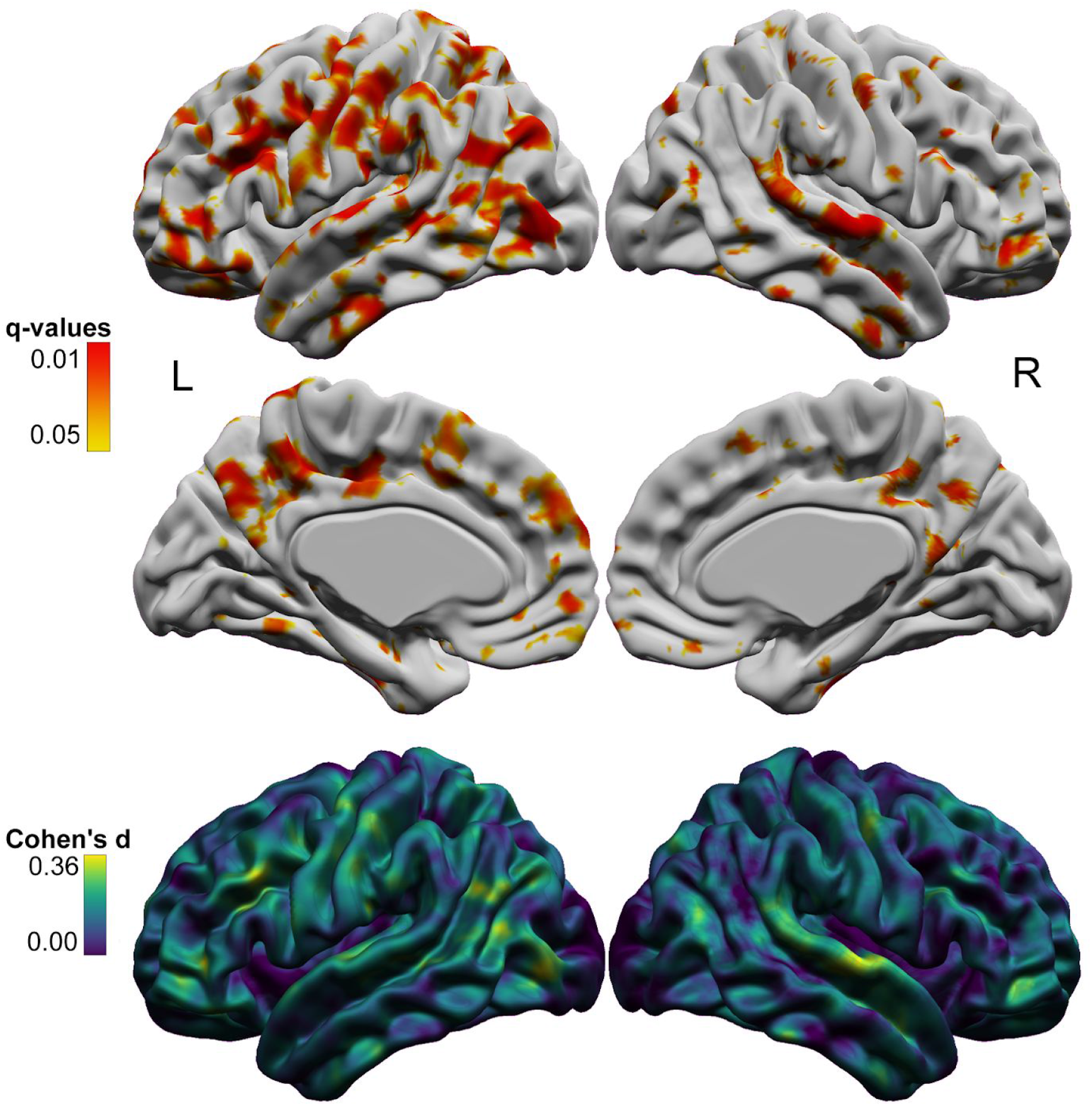
Main effect of diagnosis on BSC assessed across the entire cohort, with a smoothing kernel of 10mm (instead of 20mm). The top 4 brains display areas of significantly greater BSC in individuals with ASD 5% FDR, where the top row is showing a lateral view, and the bottom row displays a medial row (midline vertices have been removed). The bottom row displays the effect size of ASD on BSC.

**Supplementary Figure S12.**
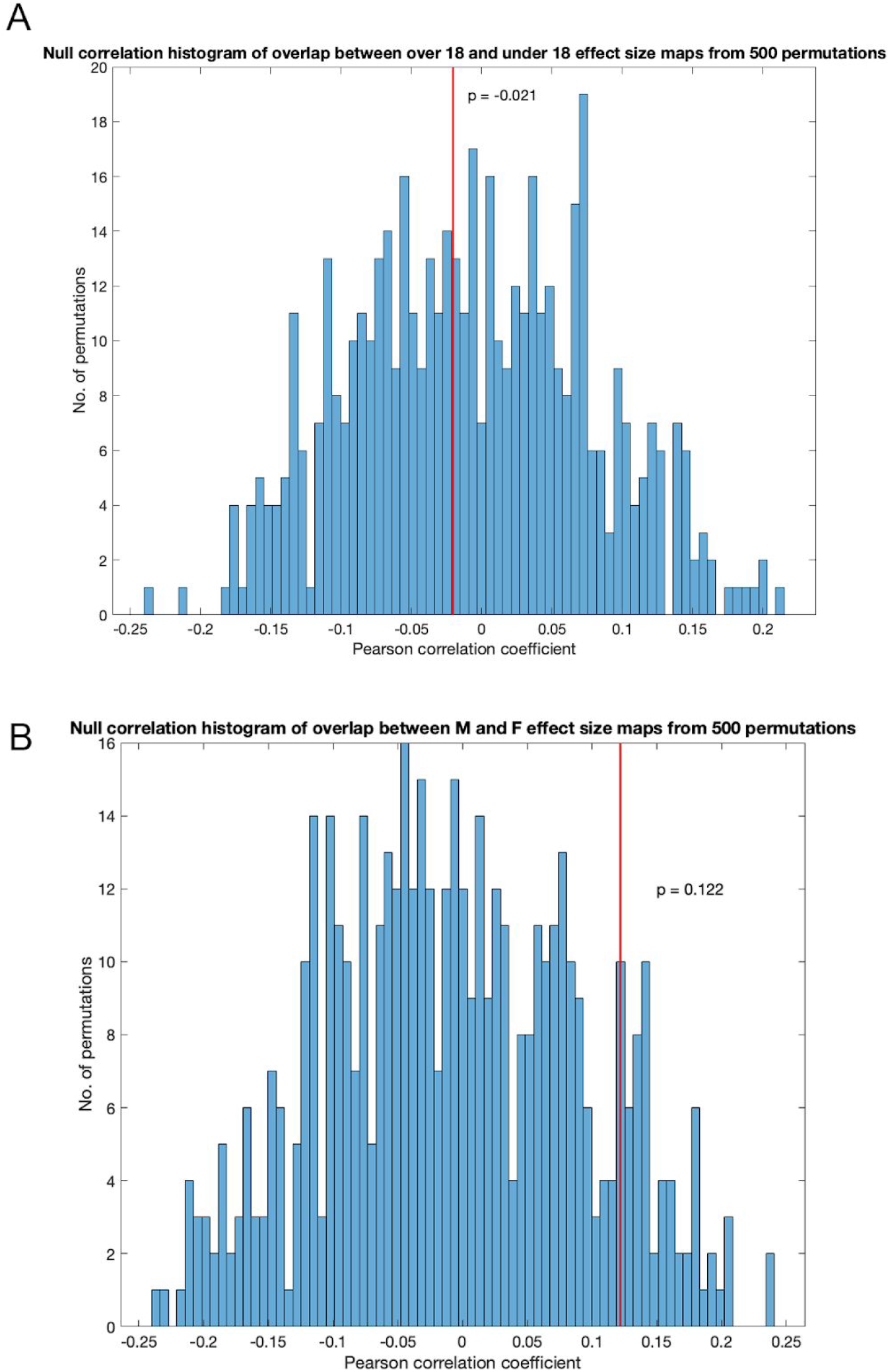
Null correlation densities from ‘spin tests’, performed between maps effect size of ASD diagnosis, in which significance of the spatial overlap of two maps is determined by rotating one map randomly, and comparing the Pearson’s correlation coefficient measured in the true overlap with those measured between the original map and rotated maps (null overlap coefficients). (A) Null correlation density from overlap between maps of effect of ASD in individuals over 18 versus those under 18, which is not significant (Rho =-0.021, p > 0.05). (B) Null correlation density from overlap between maps of effect of ASD in males and females, which is not significant (Rho=0.122. p>0.05).

**Supplementary Figure S13.**
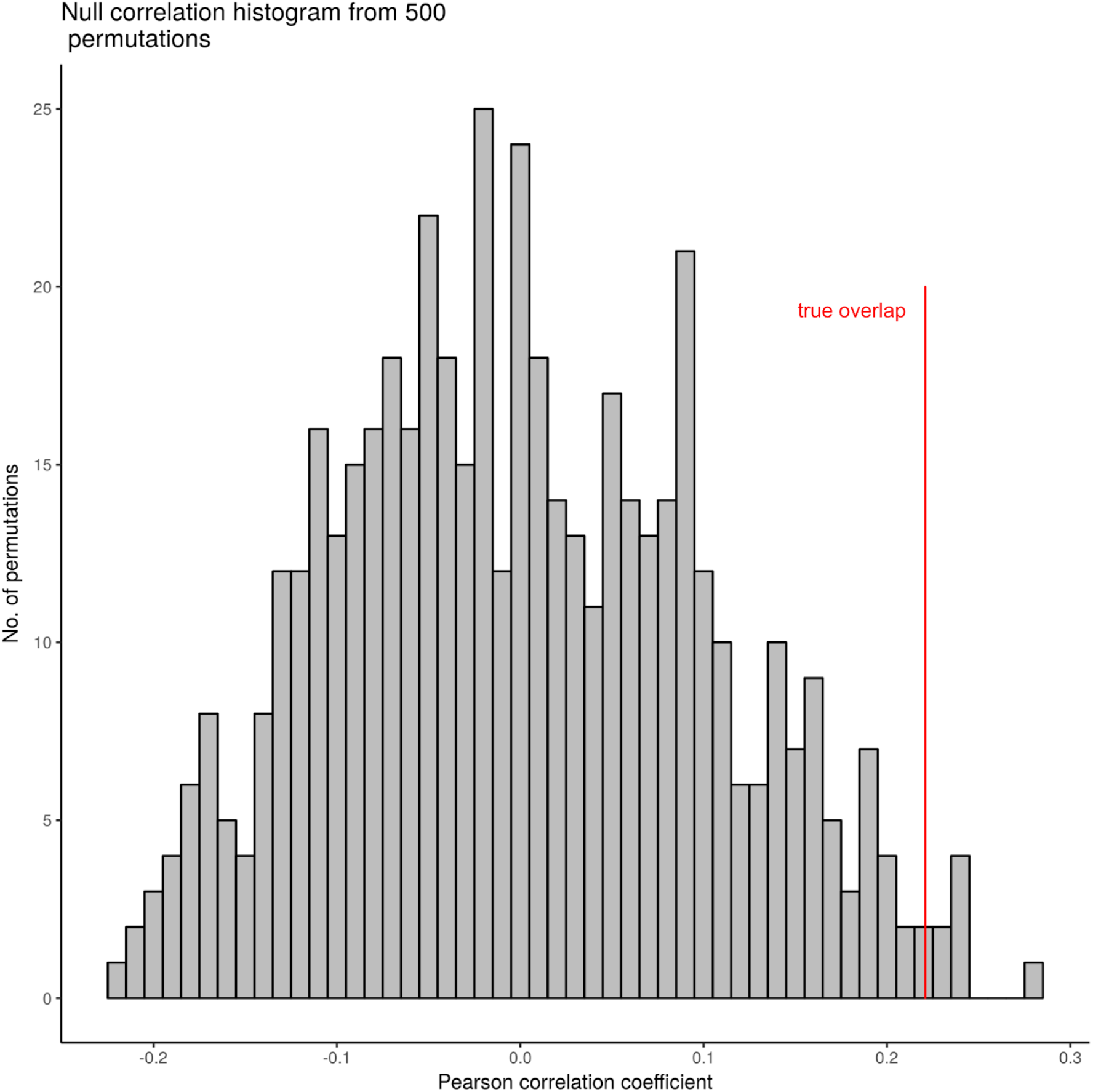
Null correlation density from overlap between maps of cortical thickness and BSC FDR-corrected q-values of main effect of ASD diagnosis (p = 0.02) generated by rotating a cortical surface randomly and comparing the Pearson’s correlation coefficient measured in the true overlap with those measured between the null models of overlap.

**Supplementary Figure S14.**
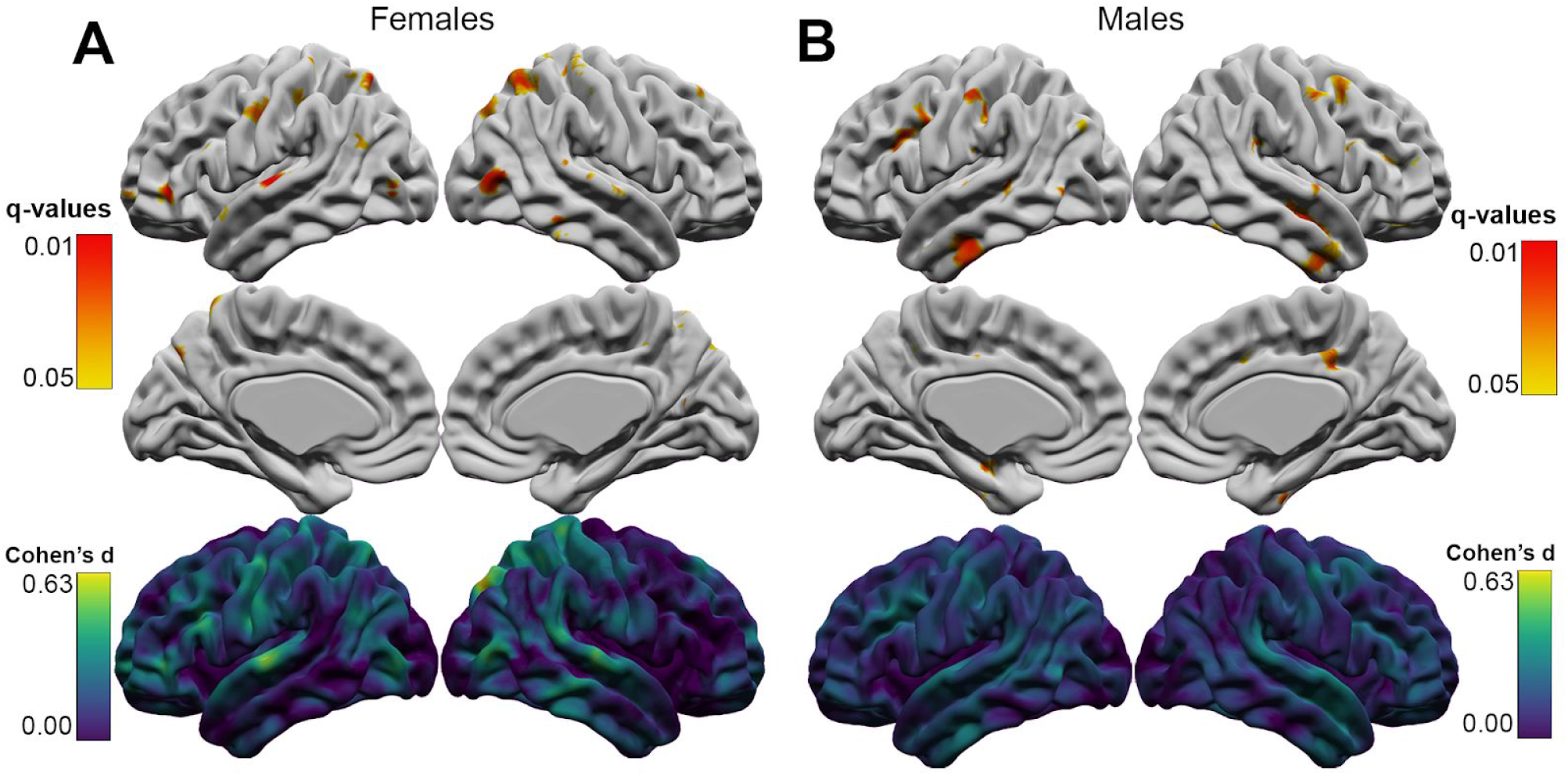
Sex-specific patterns of group differences in BSC were observed, with females showing significantly increased BSC 5% FDR in the bilateral superior parietal gyrus and superior temporal gyrus (A), and males displaying significantly greater BSC in the bilateral inferior temporal gyrus and left inferior frontal lobe (B). Further, the magnitude of the effect size was greater in females, where the peak Cohen’s d value observed in females was 0.63, and in males, it was 0.32.

## Acknowledgements

This study was performed by EO while at the Cerebral Imaging Centre, Douglas Mental Health University Institute, Montreal, Canada. E.O. is supported by the Natural Science and Engineering Research Council of Canada (NSERC) and the Fonds de la recherches en santé du Québec (FRQS).

The Cambridge Family Study of Autism was funded by a Clinician Scientist Fellowship from the MRC (G0701919) to MDS.

This study was funded by the MRC UK as the Autism Imaging Multicentre Study (AIMS), grant number GO 400061 to PIs Murphy, Bullmore, Baron-Cohen.

The Toronto cohorts were funded by the Canadian Institutes of Health Research (CIHR) MOP-106582, MOP119541 and MOP-142379 to Taylor and Anagnostou GAD is supported in part by funding provided by Brain Canada, in partnership with Health Canada, for the Canadian Open Neuroscience Platform initiative.

M.V.L. is supported by a European Research Council (ERC) Starting Grant (ERC-2017-STG; AUTISMS; no. 755816)

A.R. is supported by the intramural program of the National Institutes of Mental Health (Clinical trial NCT00001246, clinicaltrials.gov; NIH Annual Report Number, 1ZIAMH002949-03).

M.M.C is supported by Canadian Institutes of Health Research (CIHR), Natural Science and Engineering Research Council of Canada (NSERC), Fonds de la recherches en santé du Québec (FRQS), the Weston Brain Institute, and the Healthy Brains for Healthy Lives Initiative (Canada First Research Excellence Fund - McGill University).

## Notes

### Competing Interest Statement

The authors have declared no competing interest.

### Summary of Updates

In this revision, we have made several substantial changes that we believe have improved the quality of the study. In particular, we have examined normative effects of age and sex on BSC in a subset of healthy controls in order to contextualize diagnostic group differences; we have examined the relationship between BSC and estimates of in-scanner motion; and we have made the code for generating BSC publicly available on GitHub.

